# The complexity and commonness of the two-process model of sleep regulation from a mathematical perspective

**DOI:** 10.1101/2025.01.22.634299

**Authors:** Anne C. Skeldon, Derk-Jan Dijk

## Abstract

The two-process model (2pm) of sleep regulation is a conceptual framework and consists of mathematical equations. It shares similarities with models for cardiac, respiratory and neuronal rhythms and falls within the wider class of coupled oscillator models. The 2pm is related to neuronal mutual inhibition models of sleep-wake regulation. The mathematical structure of the 2pm, in which the sleep-wake cycle is entrained to the circadian pacemaker, explains sleep patterns in the absence of 24-h time cues, in different species and in early childhood. Extending the 2pm with a process describing the response of the circadian pacemaker to light creates a hierarchical entrainment system with feedback which permits quantitative modelling of the effect of self-selected light on sleep and circadian timing. The extended 2pm provides new interpretations of sleep phenotypes and provides quantitative predictions of effects of sleep and light interventions to support sleep and circadian alignment in individuals, including those with neurodegenerative disorders.

## Introduction

One of the most widely reproduced figures in sleep and circadian science must be Fig. 4 from Alexander Borbély’s 1982 paper [1] that introduces the two-process model (2pm) of sleep regulation, re-plotted in Fig. 1a. Such is the strength of the idea that two processes, i.e. sleep homeostasis and circadian rhythmicity regulate sleep timing that many data continue to be interpreted within this conceptual framework. In the 1982 paper, this conceptual framework is presented as two processes which together capture sleep propensity and duration of recovery sleep. Subsequently, in 1984, Daan, Beersma & Borbély [2] introduced twin thresholds, one determining spontaneous sleep onset and one determining spontaneous sleep offset. The enduring appeal of the powerful two-process framework is perhaps in part due to the easily interpretable visual framing of a homeostatic sleep pressure that accumulates during wake and dissipates during sleep, see Fig. 1b. The separation into circadian and homeostatic components has repeatedly been shown to provide novel insight in sleep-wake regulation, for example [3, 4].

**Figure 1.**
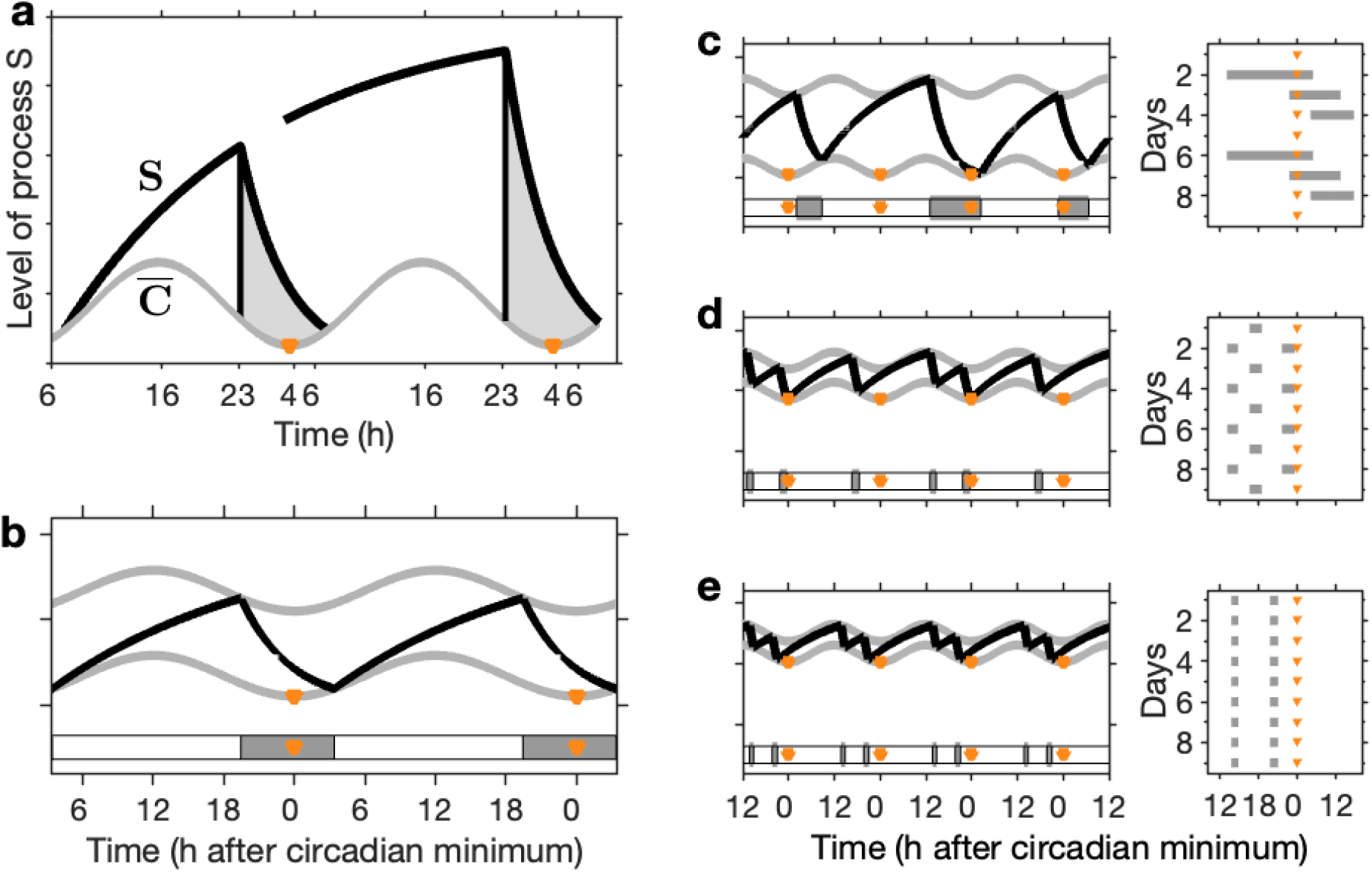
The two process model. **a** The two processes in the two-process model (2pm) as depicted in Fig. 4 from [1]. S represents the homeostatic sleep pressure, *C* is the circadian drive for wakefulness. The grey shaded region highlights times when sleep occurs. **b** The 2pm including upper and lower thresholds, as described in [2]. Homeostatic sleep pressure (black line) increases during wake and decreases during sleep. Switching between wake and sleep and sleep and wake occurs at upper and lower thresholds respectively (grey lines). The thresholds are modulated according to a circadian rhythm. Here the circadian rhythm is assumed to be entrained to 24 hours. The timing of sleep is shown by the horizontal grey bars with the orange triangles indicating the times of the circadian minimum. Parameter values: *χ*_*s*_ = 4.2 h, *χ*_*w*_ = 18.2 h, 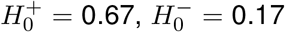, *a* = 0.12, *µ* = 1. **c–e** Simulations for different values of the mean lower threshold and *χ*_*s*_ = 4.2 h, *χ*_*w*_ = 18.2 h, 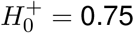, *a* = 0.07, *µ* = 1. In each pair of panels the lefthand panel shows the 2pm and the righthand panel shows a raster plot indicating the timing of sleep relative to the circadian minima (orange triangles). **c** Shows a repeating rhythm consisting of three sleep periods every four days, 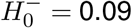. **d** Shows a repeating rhythm consisting of three sleep periods every two days, 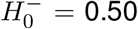. **e** Shows a polyphasic sleep pattern consisting of two sleep periods every day, 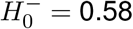.

Although often represented graphically, the 2pm consists of mathematical equations with a precise meaning. For example, the model for the sleep process, process S, can be viewed as a kinetic model in which the rate of change of homeostatic sleep pressure (be it chemical or neuronal network connectivity related processes) is a balance between the rate of increase (gain) and rate of dissipation (loss). Rate of dissipation is modelled as proportional to the instantaneous sleep pressure with rate constants 1*/χ*_*s*_ h^*−*1^ and 1*/χ*_*w*_ h^*−*1^ during sleep and wake respectively and rate of increase is at the fixed rate of *µ*_*s*_*/χ*_*s*_ (units of homeostatic sleep pressure) h^*−*1^ during sleep and *µ*_*w*_*/χ*_*w*_ (units of homeostatic sleep pressure) h^*−*1^ during wake. During wake, the rate of increase is greater than the rate of dissipation so homeostatic sleep pressure increases. During sleep, the rate of dissipation is greater than the rate of increase, so homeostatic sleep pressure decreases.(An alternative view is that there is no increase of homeostatic sleep pressure during sleep, only loss. This is equivalent to measuring sleep pressure relative to a baseline in which gain and loss are balanced during sleep.)

The mathematical equations for the 2pm include a total of eight parameters: the time constants *χ*_*s*_, *χ*_*w*_ related to dissipation and increase respectively; the ‘increase’ parameters *µ*_*w*_ and *µ*_*s*_, also known as the upper and lower asymptotes; the mean levels of the upper (wake) threshold and lower (sleep) thresholds 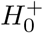 and 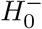 and the circadian amplitude *a* and period *T*_*c*_. Daan, Beersma & Borbély considered non-dimensional versions of the equations which effectively sets *µ*_*s*_ to zero and *µ*_*w*_ ≡ *µ* to 1. The full equations and a discussion of the scaling is given in the Methods.

The fact that the 2pm is not only a conceptual framework but also a mathematical model means it has quantitative predictive power. Indeed, Daan, Beersma & Borbély [2], showed that their model could explain sleep duration after sleep deprivation, internal desynchrony seen in time isolation experiments and enhancement and fragmentation of sleep in continuous bed rest. Following Borbély and Wirz-Justice [5], Beersma, Daan & Van den Hoofdakker [6] used the 2pm to argue that changes in sleep in depression could be explained by a reduced rate of increase of homeostatic sleep pressure during wake or increased noise in the wake threshold. Subsequently, extensive evaluation and modelling related to the increase and dissipation of homeostatic sleep pressure, as indexed by slow wake activity, has been carried out in humans (e.g. [7]), rats (e.g. [8, 9]).

A key element of the 1982 paper is the use of slow wave activity (SWA, EEG power density in the 0.75-4.5 Hz range during NREM sleep) as a biomarker for sleep pressure. However, one potential criticism is that no physiological correlates of the levels of the thresholds or the circadian amplitude have yet been found. The precise shape of the threshold, mean values and amplitude of the circadian rhythm were selected to replicate patterns of human sleep. Daan, Beersma & Borbély postulate that the upper and lower thresholds reflect circadian variation in the sensitivity of (unspecified) brain regions involved in the generation of sleep and wakefulness to homeostatic sleep pressure. In general, they argue that the upper threshold in particular may be labile and respond to short term environmental, physiological and psychological changes such as light, stimulants (coffee) and motivational aspects. Subsequent work has shown that it is possible to relate the 2pm to mechanistic neuronal models in which sleep and wake promoting neuronal populations were specified. This neuronal perspective gives further insight into possible physiological interpretations of the 2pm [10].

The 1984 paper [2] highlights the rich dynamics of the 2pm and its ability to simulate the dependency of sleep duration on circadian phase of sleep onset but also the appearance of both monophasic and polyphasic sleep, even though the period of the circadian process remains within the 24-h range, see also Fig. 1c-e. From a mathematical perspective it is natural to ask what the underlying structure of the model is that leads to these different phenomena, and then what does that mean for explaining data. In fact, around the time the 2pm was formulated, similar ‘threshold’ models were constructed to explain phenomena such as integrate-and-fire patterns in neuronal systems [11, 12], breathing (inspiration-expiration) phenomena [13, 14], and cardiac arhythmias [15, 16], see Fig. 2. All these models explaining phenomena across a wide range of time scales have the same underlying mathematical structures, which can be formally analysed via circle maps [17]. Circle maps are mathematical objects which have been studied in their own right [18] and for which many statements on the existence and type of dynamical behaviour have been proved. Using the circle map representation means that the dynamics of different threshold models can be brought into a unifying framework and understood via established theorems. In Fig. 2, we also show the associated circle maps for each of the example models. An explanation of how circle maps are constructed is given in the Supplementary Fig. S1.

**Figure 2.**
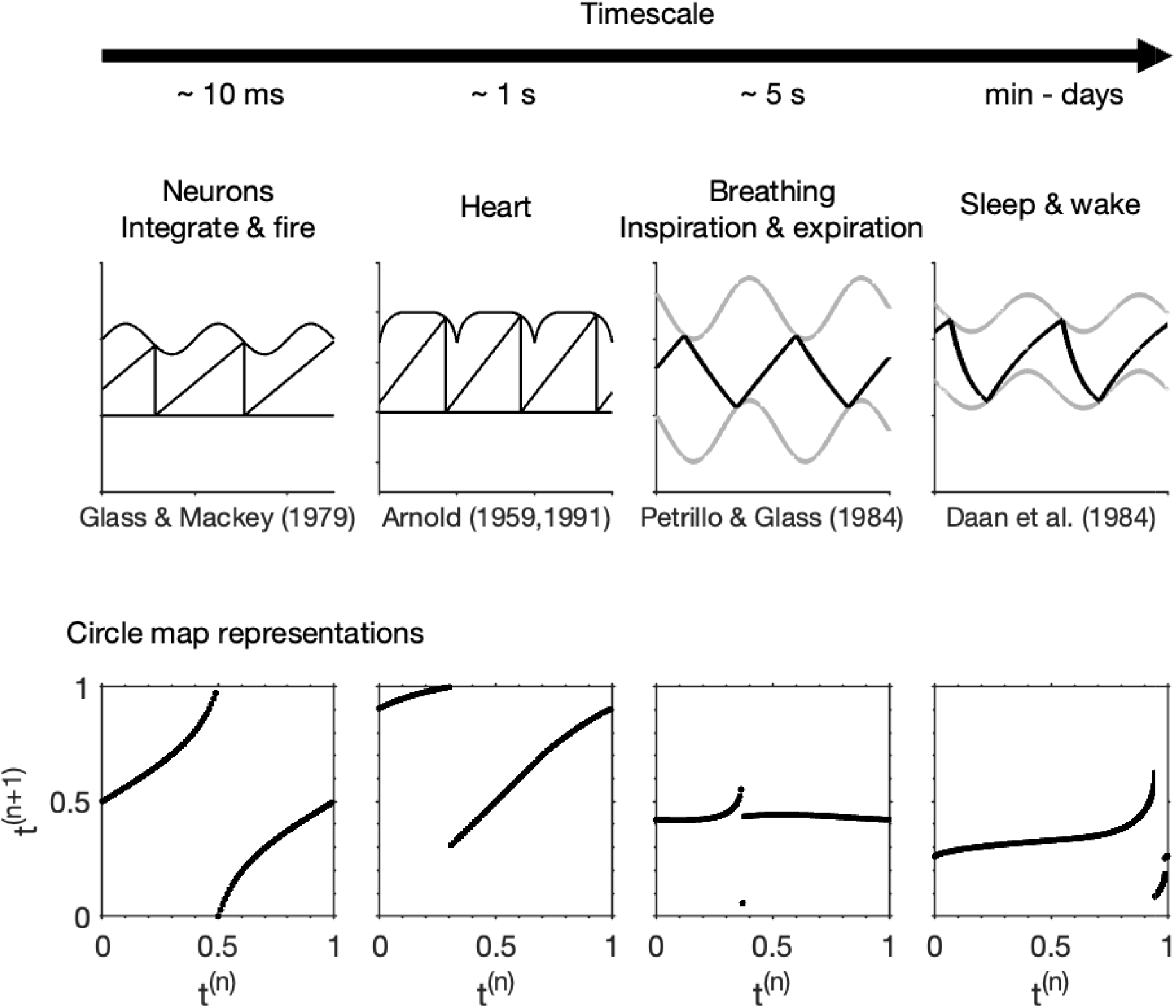
Threshold models. Four different threshold models for physiological processes acting on different timescales including neuronal [11], cardiac [15, 16], breathing [14] and sleep-wake regulation [2]. Each model can be represented either as a timeseries, as in the upper panels, or a circle map, as illustrated in the lower panels. The circle map description is constructed by using the fact that each threshold model is essentially a rule that describes how one time of hitting the upper threshold maps to the next time of hitting the upper thresholds. Since in all models, the threshold is periodic with (scaled) period 1, this rule can be considered as a rule that takes one value in the interval [0, 1) and maps it to another value in the interval [0, 1). Further detail on the circle map construction is given in Supplementary Fig. S1.

The mathematical perspective highlights that the 2pm sits within the general class of systems which consists of an oscillation with a so-called natural period (the sleep oscillator) which is forced by a periodic input (the circadian oscillator) [19]. All the structures that are standardly seen in such systems, such as tongue-like regions of entrainment describing the phase relationships between the two oscillators and the number of oscillations observed per period of the forcing oscillator, are then seen in the 2pm.

The 2pm describes the duration of sleep and the timing of sleep relative to the phase of the circadian oscillator for which the circadian minimum is often assigned phase 0. When used to describe timing relative to the 24 hour day two important assumptions are made. First, that the circadian period *T*_*c*_ is entrained to 24 hours, so the periodicity of the circadian forcing *T*_*f*_ is 24 hours. Second, that the circadian minimum occurs at a specific clock-time, e.g. approximately 06:00 in young adults [20]. However, the 2pm as described in the 1982 and 1984 papers, does not include a mechanism to explain entrainment to 24 hours, i.e. the timing of the circadian pacemaker to its external forcing oscillator which is now generally considered to be the light-dark cycle. Daan, Beersma & Borbély were aware of this limitation, but commented that there was insufficient knowledge of the phase response of the circadian oscillator of humans to the light-dark oscillation to warrant inclusion of a quantitative description of how light-dark cycles could entrain the circadian process. Consequently, in its original form, the 2pm is limited in the extent to which it can explain sleep and circadian timing relative to the light-dark cycle. However, since 1984 more has been discovered about the light-circadian entrainment process. This has led to extensions of the 2pm which include light and describe the system governing human sleep-wake timing as three interacting (coupled) oscillatory processes [21].

In this paper we briefly discuss three aspects of the 2pm: first, the relation to neuronal mutual inhibition models of sleep-wake regulation, the consequent physiological interpretation of the thresholds and the wider implications; second, the mathematical structure that explains the internal desynchrony results, the occurrence of bicircadian rhythms and polyphasic sleep, and third, extensions of the 2pm model to include interaction with the light-dark cycle. In the process, we draw together and expand on previously published work, some of which has primarily been written for mathematical audiences. We aim to close the gap between descriptions of the mathematical and physiological/behavioural aspects of the 2pm. We emphasize that recognition of the interaction between three oscillators governing sleep-wake timing in humans leads to novel interpretations of sleep phenotypes. We end with a brief discussion of opportunities and future directions based on the extended 2pm.

## Results

### Neuronal mechanisms underpining the two-process model

Since the 1980’s we have learned much about the brain structures involved in sleep regulation. Saper, Scammell & Lu [22] conceptualised sleep-wake regulation as the result of the mutual inhibition of wake active and sleep active regions in the brain. This conceptual framework explains the stability of sleep and wake states: during wakefulness, wake-active neurons fire and suppress the firing of sleep-active neurons, thereby maintaining wake. When asleep, sleep-active neurons fire, and by suppressing the firing of wake-active neurons, maintain sleep. Importantly, this mutual inhibition framework alone does not explain why switching between sleep and wake states occurs. Consequently, Saper, Scammell & Lu additionally proposed the existence of time dependent inputs to wake active and sleep active regions. These time dependent inputs reflect homeostatic sleep pressure and circadian rhythmicity. When these inputs are sufficiently strong, they overcome the inhibition and enable switching between states to occur.

The conceptual neuronal sleep-wake regulation framework was formulated in a mathematical model by Phillips and Robinson [23] with interactions as shown in Fig. 3a. Simulations of the Phillips-Robinson model capture the rise of sleep pressure during wake and decrease during sleep, the oscillations of the circadian rhythm and the so-called flip-flop between firing of sleep active and wake active neurons, see Fig. 3b-e.

**Figure 3.**
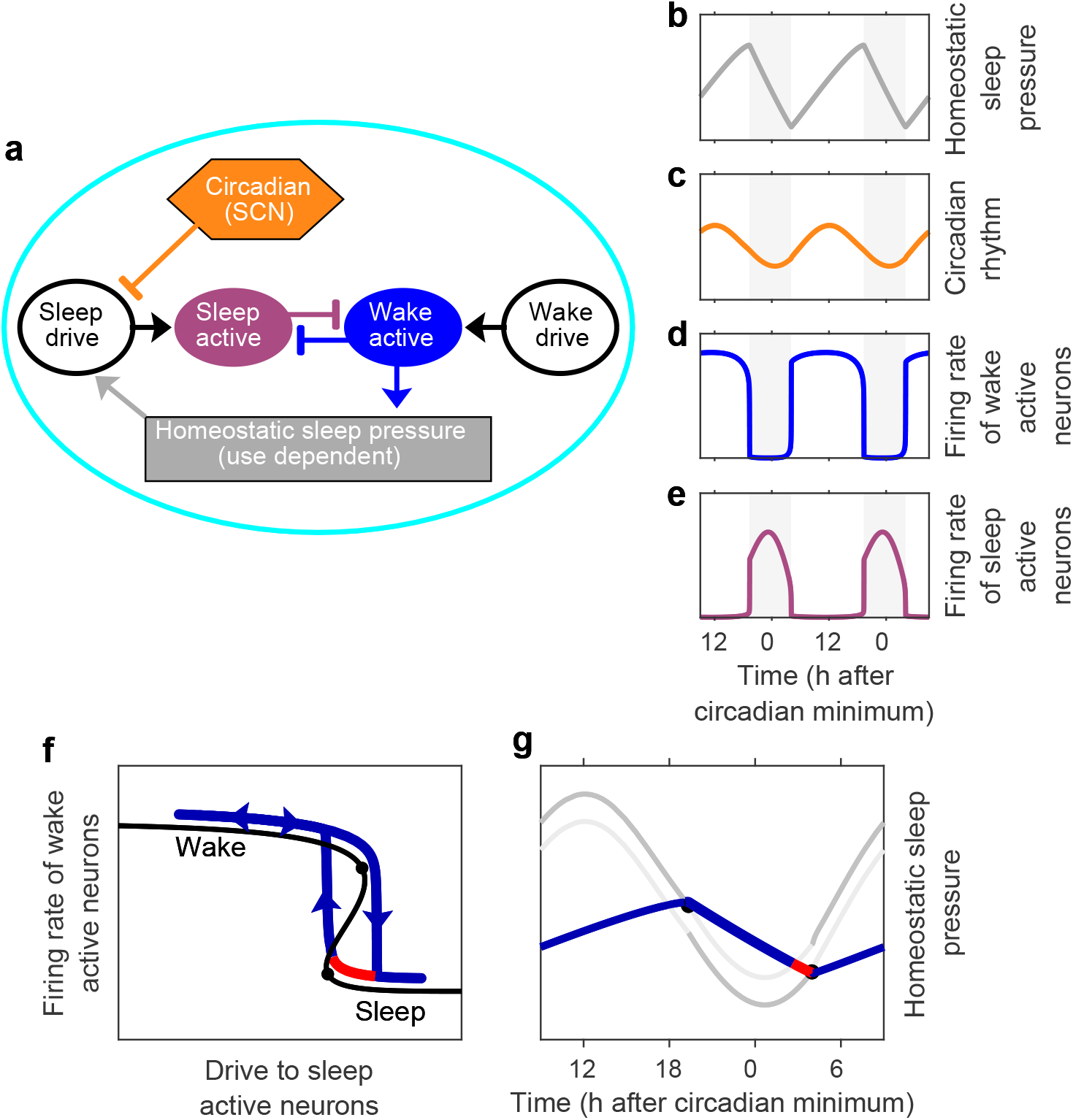
A neuronal mutual inhibition model and its relation to the two-process model. **a** Neuronal structure underpinning sleep-wake regulation, based on the conceptual framework of Saper, Scammell & Lu [22] as framed by Phillips & Robinson [23]. (In this model the impact of the suprachiasmatic nucleus (SCN) is primarily by variation of inhibition of the sleep drive. Based on physiology, it is more likely that the SCN primarily activates the wake promoting part of the system during most of the 24 h day, with only small contributions to sleep drive late in the biological night [25]). **b**–**e** Simulation of the Phillips-Robinson model for standard parameter values. **b** Homeostatic sleep pressure increases during wake and decreases during sleep. **c** Sinusoidal representation of the circadian rhythm from the SCN entrained to 24 h. **d** Firing rate of wake active neurons (high during wake and low during sleep). **e** Firing rate of sleep active neurons (high during sleep and low during wake). **f** Hysteretic representation of the Phillips-Robinson model. In this representation, there is a folded surface which results from the mutual inhibition of the wake active and sleep active neuronal populations. The time dependent sleep drive results in a trajectory (blue/red) that evolves over the surface, switching between the wake (upper) surface to the sleep (lower) surface at the folds (tipping points) which are indicated by the black circles on the black line. **g** The Phillips-Robinson model in the form of the two-process model. The homeostatic sleep pressure (blue / red curve) increases during wake and decreases during sleep and is colour-coded as in panel **f**. The mean level of the upper (lower) threshold is given by the value of the drive at the upper (lower) fold in panel **f**. The oscillation of the thresholds is given by the circadian inhibitory input to the sleep drive. The hysteretic structure suggests that it is the upper threshold that is of prime relevance during wake, and the lower threshold which is of prime relevance during sleep, we have therefore drawn the relevant ‘active’ threshold in dark grey and the ‘inactive’ threshold in light grey.

**Figure 4.**
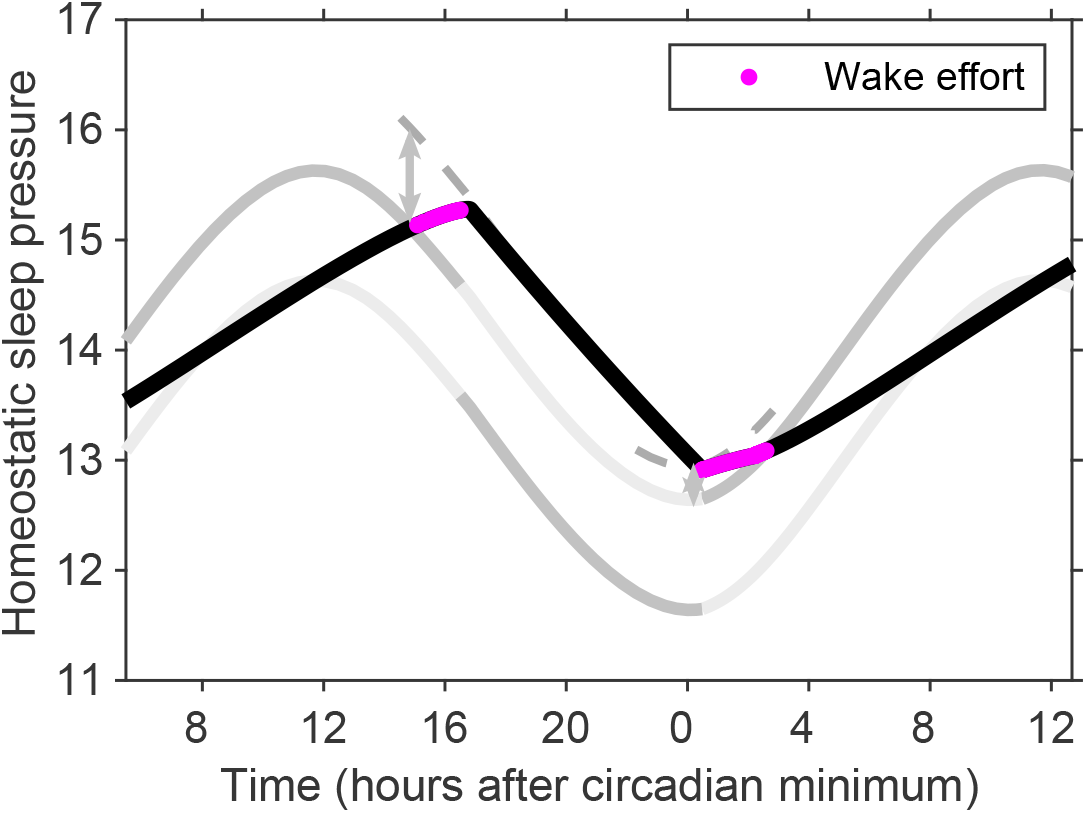
Wake effort. In the neuronal interpretation of the two-process model, during wake, once the upper threshold has been reached requires effort i.e. additional input to wake active neurons which shifts the position of the upper threshold. If awoken while homeostatic sleep pressure is above the upper threshold, then again effort is required to shift the upper threshold (grey dashed lines). Regions in which wake effort is needed are highlighted in magenta. As in Fig. 3 we have drawn the ‘active’ parts of the thresholds in dark grey and the ‘inactive’ parts of the thresholds in light grey.

The Phillips-Robinson model has two timescales. A fast timescale, governing neuronal dynamics on the order of seconds, and a slow timescale, regulating sleep homeostasis and circadian rhythmicity over hours. This results in simulated sleep-wake cycles in which typically there are rapid transitions between sleep and wake, but that the states of sleep and wake last for hours, as in Fig. 3d,e. This multiple timescale structure means that an alternative representation of the Phillips-Robinson model is as a hysteresis loop, with transitions between wake and sleep and sleep and wake occurring at tipping points, see Fig. 3f, [23, 24]. Furthermore, using a formal mathematical multiple timescale analysis, on the slow timescale the Phillips-Robinson model may be mapped to the 2pm [10]. In this mapping, the tipping points that occur in the neuronal model are equivalent to the switching points between wake and sleep in the 2pm.

Often the 2pm is described as homeostatic sleep pressure oscillating ‘between’ two thresholds, but the neuronal framework highlights that requiring sleep pressure to be between the thresholds is not a requirement for the model to successfully explain patterns of sleep behaviour. For example, Fig. 1b and Fig. 3g have the same sleep duration and relation between sleep and circadian timing but the gap between the thresholds in Fig. 3g is much smaller and the circadian amplitude much lower than in Fig.1b. In fact, the hysteretic structure suggests that at any one point, only one of the thresholds is of relevance i.e. the lower threshold when asleep (on the lower branch of the hysteresis loop) and the upper threshold when awake (on the upper branch of the hysteresis loop). To highlight the fact that at any one time only one threshold is relevant, in Fig. 3 we have distinguished between the ‘active’ and the ‘inactive’ threshold. This raises interesting interpretative differences. Based on a view that sleep and wake correspond to a hysteretic structure, the region between the thresholds is a region in which both sleep and wake states exist. Above the upper threshold, only the state of sleep can exist and below the lower threshold, only the state of wake can exist. The hysteretic structure suggests that the Phillips-Robinson model, and by extension the two-process model may be alternatively viewed via a potential formulation, see Supplementary Fig. S2.

This hysteretic structure has at least three consequences. First, that between the two thresholds, switching between sleep and wake may occur through random perturbations. Second, when homeostatic sleep pressure reaches the upper threshold, either sleep ensues or some additional wake ‘effort’ is required to shift the upper threshold. Third, if woken while still in the ‘sleep only’ region, then to remain awake requires additional wake drive to shift the upper threshold. This has been conceptualised as ‘wake effort’ [26, 27]. In modelling acute sleep deprivation in humans, Daan, Beersma & Borbély viewed that the upper threshold was over-ridden. In the wake effort view, the upper threshold is moved to allow wake to be maintained, as illustrated in Fig. 4. The fact that sleep pressure does not remain between the thresholds in the Phillips-Robinson parameter regime has implications for waking early. In the Daan, Beersma & Borbély parameter regime, waking early requires momentary stimulation to switch from one stable state (sleep) to the other stable state (wake). In the Phillips-Robinson parameter regime, wake can only be maintained by ‘wake effort’ shifting the position of the upper threshold. The requirement of additional effort to maintain wake remains until late enough in the day that the habitual rise in circadian drive for wakefulness means that wake is a stable state, see Fig. 4.

The correspondence between the 2pm and Saper’s framework, via the Phillips-Robinson model, means that many of the results that have been found from simulating the Phillips-Robinson model can also be seen in simulations of the 2pm and vice versa. Furthermore, there is a direct relationship between the parameters in the neuronal model and the levels of the thresholds in the 2pm. For example, increasing the drive to wake active regions predominantly has the effect of raising the level of the upper threshold. Increasing the drive to sleep active regions lowers the mean value of the thresholds while leaving the distance between then unchanged, see Fig. 5. The positions of the thresholds are additionally affected by parameters that relate to the strength of the mutual inhibition and the neuronal firing rates, see Supplementary Fig. S3.

**Figure 5.**
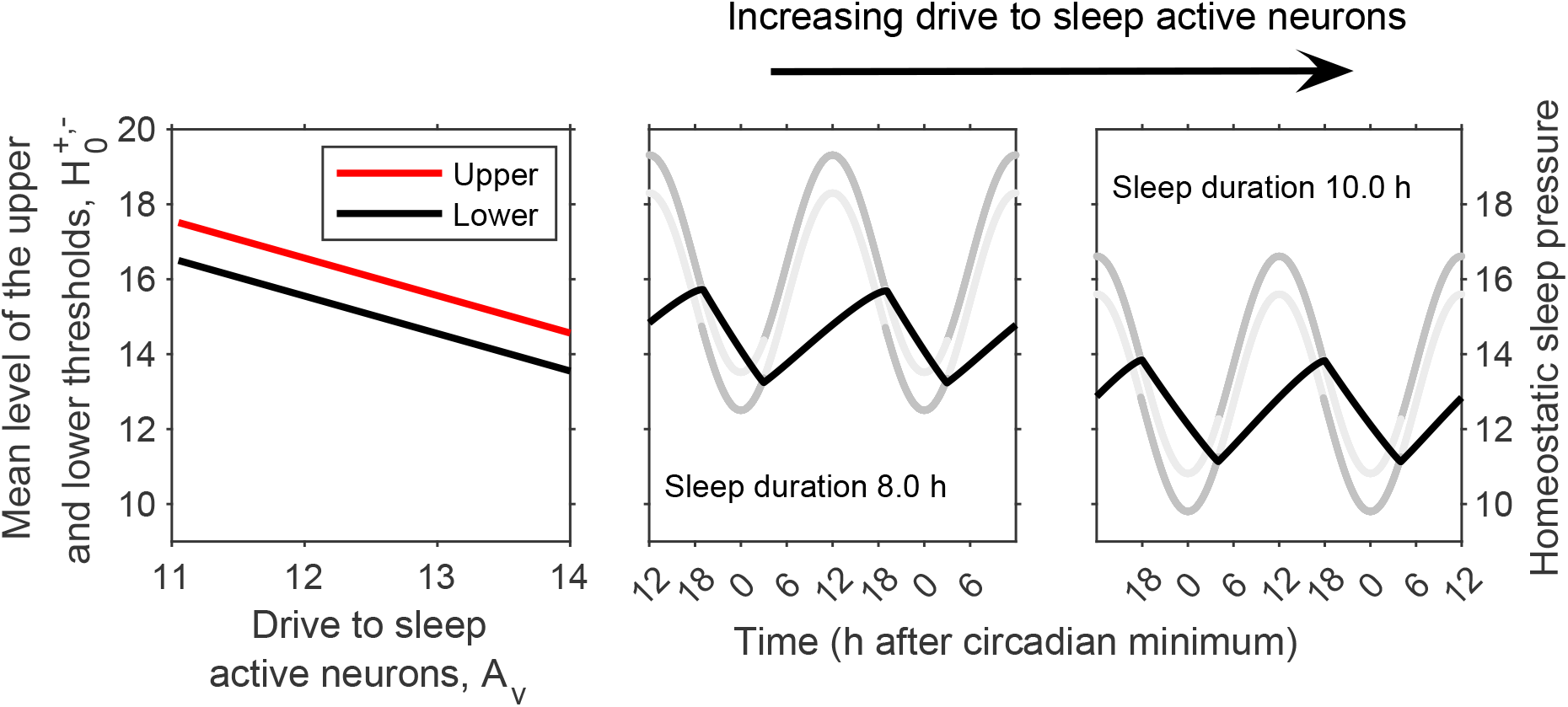
Increasing drive to sleep active neurons decreases the mean levels of the thresholds. Simulation of the Phillips-Robinson model [23] for different values of the drive to sleep active neurons. The gap between the thresholds in the 2pm model do not change, but lowering the thresholds increases sleep duration because of the exponential nature of the model for homeostatic sleep pressure. As in Fig. 3 we have drawn the ‘active’ parts of the thresholds in dark grey and the ‘inactive’ parts of the thresholds in light grey.

### The mathematical structure of the two-process model

#### The sleep-wake oscillator

In the absence of circadian rhythmicity (zero circadian amplitude), the 2pm reduces to a homeostatic sleep pressure oscillating between time-independent upper and lower thresholds, see Fig. 6a. This sleep-wake oscillator is also often described as a relaxation oscillator. The natural period *T*_nat_ of the oscillation can be formally derived in terms of parameters of the two-process model and is given by

**Figure 6.**
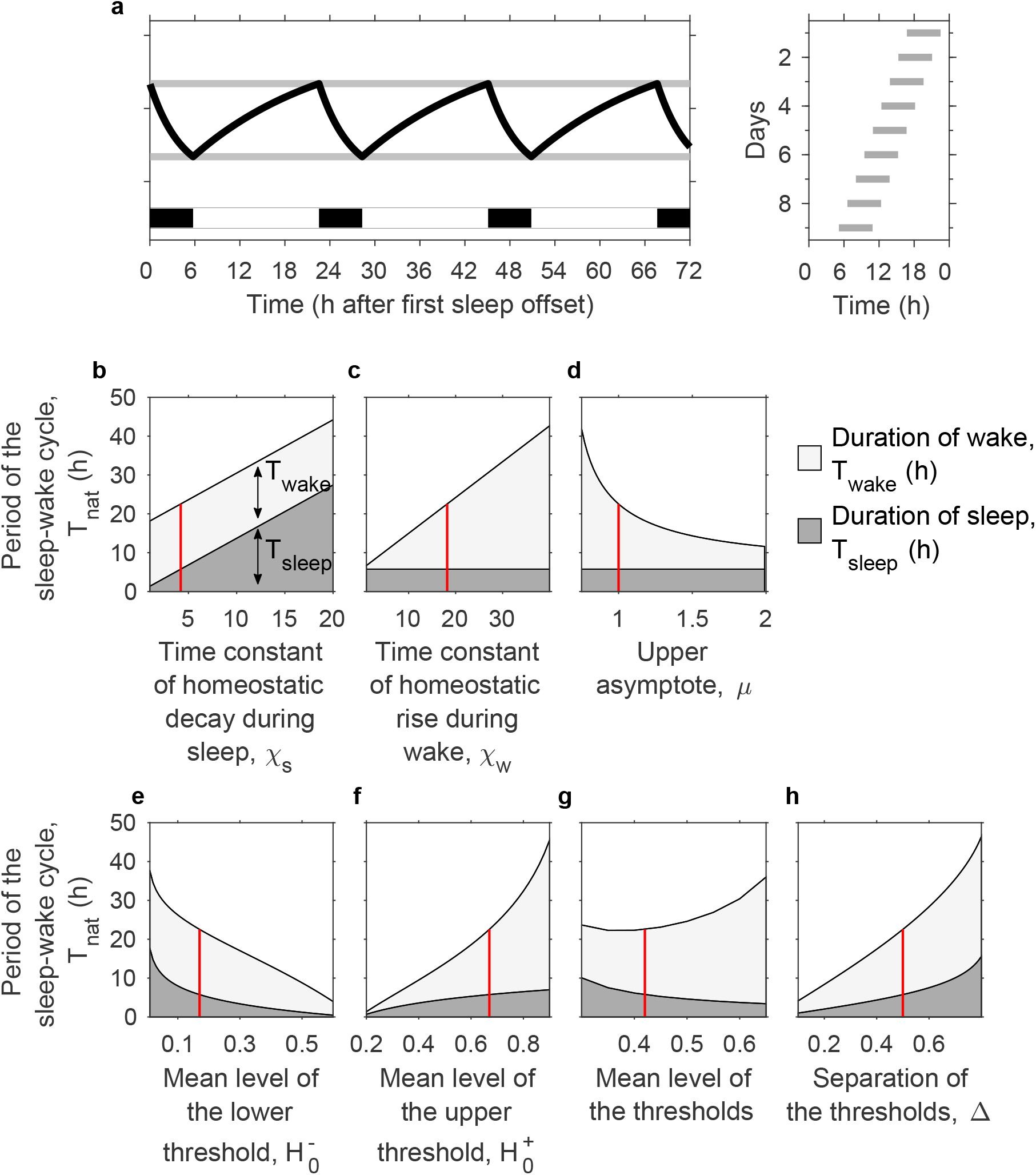
Natural period of the sleep-wake oscillator. **a** The two-process model (2pm) with the circadian amplitude set to zero. For the standard parameters (*χ*_*s*_ = 4.2 h, *χ*_*w*_ = 18.2 h, 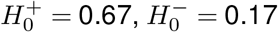, *µ* = 1) the period of the sleep-wake cycle is approximately 22.6 h, of which approximately 5.8 h is sleep (26%). The righthand panel shows a raster plot with the timing of sleep on successive 24 h days. **b-h** Illustration of the dependence of the natural period on the parameters in the 2pm, as given by equations (1)-(3). The vertical red line indicates the position of the standard parameters. Since it is sometimes useful to describe the position of the thresholds by a mean value and the separation instead of an upper and lower value, we have shown the dependence of sleep and wake duration on the value of the upper threshold 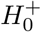, the value of the lower threshold 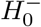 and also the mean level of the thresholds 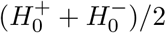 and the separation of the thresholds 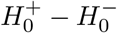.

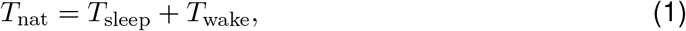

where *T*_sleep_ is the duration of each sleep period and *T*_wake_ is the duration of each wake period. Here,

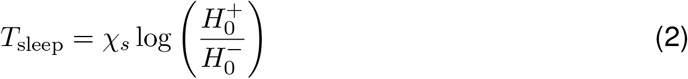

and

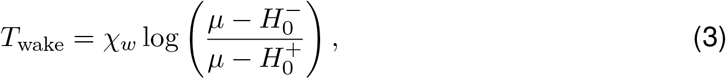

where *χ*_*s*_ and *χ*_*w*_ are the homeostatic time constants for sleep and wake respectively, 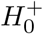 and 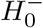 are the mean positions of the upper and lower thresholds and *µ* is the upper asymptote. The derivation of equations (2) and (3) is given in the Methods. For the standard parameters, *T*_sleep_ = 5.8 h, *T*_wake_ = 16.8 h, giving a natural period *T*_nat_ = 22.6 h in which approximately 26% is spent asleep.

The dependence of *T*_nat_, *T*_sleep_ and *T*_wake_ on the parameters is shown in Fig. 6b-h. In line with the graphical and computational intuition described in Daan, Beersma & Borbély [2], *χ*_*s*_ changes sleep duration but has no impact on wake duration, *χ*_*w*_ and *µ* change wake duration but have no impact on sleep duration. The exponential nature of sleep homeostatic pressure, as modelled, means that the positions of the thresholds 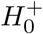 and 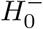 alter both sleep and wake duration. We note that the mean level of the thresholds also impact the average level of sleep pressure and the effects of sleep deprivation.

### Entrainment of the sleep-wake oscillator by the circadian oscillator

In the presence of circadian rhythmicity (circadian amplitude *a >* 0), the 2pm consists of a sleep-wake oscillator that is forced by the circadian oscillator. Characteristic of nonlinear forced oscillator systems, if the natural period *T*_nat_ of the sleep-wake oscillator is sufficiently close to the forcing period *T*_*f*_, and the amplitude of the forcing is sufficiently high then entrained monophasic sleep-wake cycles of period *T*_*f*_ result. If the forcing is weak (i.e. low circadian amplitude) then only a small range of natural periods *T*_nat_ can be entrained to *T*_*f*_. The stronger the forcing, the larger the range of natural periods can be entrained. This means that, in a (natural period)–(forcing amplitude) parameter plane, regions where entrained sleep-wake patterns consisting of one sleep-wake cycle occurs in every circadian period (monophasic sleep), form a tongue-like region, for example see Fig. 7a. We note that tongue-like entrainment regions are typical of forced systems of self-sustained oscillators: a similar tongue-like entrainment structure is seen in models of the circadian system forced by the light-dark cycle [28].

**Figure 7.**
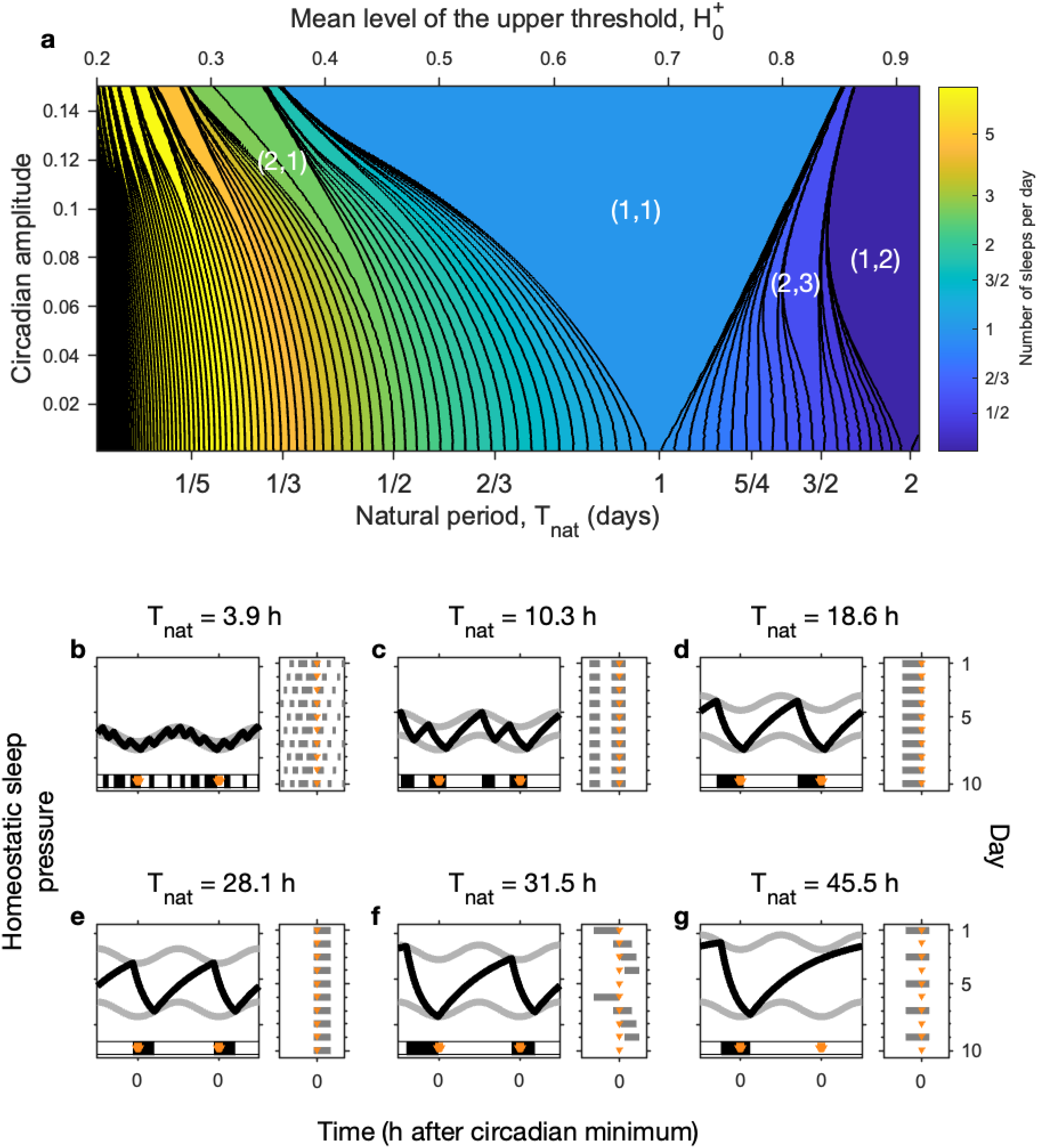
Entrainment of the sleep-wake oscillator by the circadian oscillator: Arnold tongues. **a** Number of sleeps per day as a function of the mean level of the upper threshold, 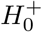 and the circadian amplitude, generated by simulating the 2pm for approximately 110,000 different parameter values. The remaining parameters are: *χ*_*s*_ = 4.2 h, *χ*_*w*_ = 18.2 h, 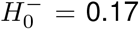, *µ* = 1. Regions are shaded according to the number of sleeps per day. The black lines are contour lines. Both the shading and the contours are on a logarithmic scale (base 2). The largest region corresponds to monophasic sleep. In principle there are regions of *n* sleeps per day for *m* circadian periods for every integer pair (*n, m*) for every *n/m*. In practice, the upper and lower limits of the values for *n/m* are bounded by the choices of the other parameters. For the example shown 0.068*< n/m <* 2.54 days. Each region forms a tongue with the tip of the tongue occurring at zero circadian amplitude for the value of 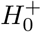 which gives a natural period of *m/n*. **b-g** Examples of simulations for different values of 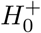 at a fixed circadian amplitude of *a* = 0.07, i.e. taking a horizontal line across panel **a**. In each case the two-process model along with a raster plot is shown. The orange triangles mark the time of the circadian minimum. **b** Polyphasic sleep pattern. **c** Two sleeps a day. **d** Monophasic sleep with the circadian minimum occurring near the end of the night. **e** Monophasic sleep with the circadian minimum occurring near the beginning of the night. **f** Four sleeps every five days. **g** One sleep every two days.

As observed by Daan, Beersma & Borbély [2], the ostensibly ‘simple’ 2pm exhibits many other dynamical behaviours. Indeed, entrainment theory predicts that there exists an entrainment tongue in which *n* sleep-wake oscillations occur in *m* periods of the circadian drive for every integer pair (*n, m*). For example, the (2,1) tongue consists of sleep-wake cycles in which two sleeps occur in each circadian period. The (1,2) tongue consists of sleep-wake cycles in which one sleep occurs every two circadian periods. The largest tongue is the (1,1) tongue, corresponding to monophasic sleep patterns.

This mathematical structure has a very particular ordering, as shown in the example in Fig. 7a. For sufficiently small circadian amplitudes, this structure is precisely ordered, solutions are unique, complicated but not chaotic. For higher circadian amplitudes, it is to be expected that solutions become chaotic and that more than one solution may exist for the same parameter values [29, 30]. Within each tongue, the phase relationship between the sleep-wake cycle and the circadian rhythm changes. This is easiest to see in the large tongue for monophasic sleep. When the natural sleep-wake period is longer than the forcing period, the forcing has to speed up the sleep-wake oscillator, with the effect that the sleep-wake oscillator tends to be ‘behind’ the circadian oscillator in phase. Conversely, when the natural sleep-wake period is shorter than the forcing period, the forcing has to slow down the sleep-wake oscillator, with the effect that the sleep-wake oscillator tends to be ‘ahead of’ the circadian oscillator in phase. The net effect is that on the side of the tongue with the longest natural period (righthand edge for Fig. 7a) the circadian minimum occurs at the beginning of the night, see Fig. 7f. For the side of the tongue with the shortest natural period, the circadian minimum occurs at the end of the night, see Fig. 7e. The example shown in 7 comes from varying the mean level of the upper threshold. Varying the mean level of the lower threshold, the homeostatic time constants or the value of the upper asymptote similarly change the natural period. Varying any one of these parameters results in a similar set of tongues, but squashed or stretched because the relationship between the natural period *T*_nat_ and the parameters differs, see Supplementary Fig. S4.

This mathematical structure follows from entrainment theory because the 2pm is an example of a circle map [31, 10]. Circle maps were first introduced by Arnold, [15], and the dynamics of circle maps have been extensively analysed (e.g. see [18, 29, 30]) and also in the particular context of the 2pm [19]. The ‘tongue-like’ structure is shared by threshold models for other systems (e.g. [11, 12, 15, 16]), and has been formally analysed via circle maps [17]. Understanding the entrainment structure facilitates understanding and interpreting data, we discuss three examples below.

### Using the mathematical structure of the two-process model to explain patterns of sleep

#### Internal desynchrony and bicircadian rhythms in humans

Time isolation experiments in which knowledge of clock time and all access to environmental timing cues are removed have been critical in the understanding of circadian rhythmicity and sleep-wake regulation. Many experiments were carried out in purpose build facilities [32, 33, 34] and in caves [35]. Strogatz [36] summarised many of the available data and evaluated the ability of various oscillator models to explain the results. In time isolation experiments, an immediate observed effect was synchrony with the 24-hour day was lost and replaced by rhythms where sleep and circadian rhythms were in synchrony with each other but typically with a 25 hour period. The phase of the minimum of the core body temperature was observed to shift from the end of the night to the beginning of the night [33]. Subsequently, in many subjects internal desynchrony was observed, in which the circadian cycle typically became slightly shorter than 25 hours but the sleep-wake cycle desynchronised oscillating with a period that was either substantially longer or shorter than the circadian cycle. In some cases, a transition to bicircadian rhythms in which there is one sleep every two days was observed.

Considering the underlying structure of the 2pm, as illustrated in Fig. 7 provides plausible explanations for how such sleep-wake patterns may occur. The initial loss of synchrony with the 24-h day results in an immediate shift in period of the circadian oscillator from its entrained value of 24 hours to approximately 25 hours. The overall tongue-like entrainment structure remains, but with small quantitative differences, for example, the minimum of the monophasic sleep tongue occurs at 25/24 instead of one day. Indeed, a formal mathematical scaling shows that changing from a forcing period of *T*_*f*_ = 24 h to *T*_*f*_ = 25 h is equivalent to rescaling time to circadian days rather than calendar days and scaling the homeostatic time constants by a factor of 25/24 days i.e. 4%. The immediate change from *T*_*f*_ = 24 h to *T*_*f*_ = 25 h therefore results in sleep-wake patterns which are entrained to the new forcing period of 25 h, see Supplementary Fig. S5a and b.

The observed change in phase of sleep relative to the circadian pacemaker can be explained by either a reduction in circadian amplitude or by a lengthening of the natural period *T*_nat_ see Supplementary Fig. S5c.

Note that changing the circadian amplitude has minimal change in the percentage of time spent asleep. In the model, changing the natural period can either increase or decrease the percentage of time spent asleep depending on which parameter is varied. The occurrence of internal desynchrony occurs in regions close to the boundary of the monophasic sleep tongue. Formally, in the 2pm, the desynchronised patterns are generically periodic patterns of period much longer than the circadian period. For low circadian amplitude, the duration of sleep will be approximately the same every day. For higher circadian amplitudes, the duration of sleep will depend on the timing of sleep relative to the timing of the circadian rhythm resulting in occasional long sleeps, see Supplementary Fig. S5d and e. This latter phenomena is analogous to ‘relative coordination’ seen in desynchrony of circadian rhythms to the 24-hour day [37].

Finally, to explain the occurrence of bicircadian rhythms within the 2pm framework requires there to be a lengthening of the natural period of the sleep-wake oscillator *T*_nat_. Since the period is given by equation (1), lengthening of the period occurs if the homeostatic time constants increase, the upper threshold increases (see Fig. 7a and Supplementary Fig. S5g) the lower threshold decreases or the upper asymptote decreases. It is not immediately evident which parameter explains the observed phenomena, although further using the percentage of time spent in sleep could establish which may be most plausible. Daan, Beersma & Borbély primarily focussed on varying the level of the upper threshold and the circadian amplitude [2]. Subsequently, Daan *et al* highlighted that low mean rectal temperature (low metabolic rate) was associated with long sleep-wake periods and high mean rectal temperature (high metabolic rate) was associated with short sleep-wake periods [38]. For the 2pm, this would suggest that higher metabolic rate are reflected in shorter homeostatic time constants, or a decreased gap between the thresholds (lower level of the upper threshold or higher level of the lower threshold) or a higher value of the upper asymptote.

In their neuronal model, Phillips-Robinson model [39] showed that internal desynchrony and bicircadian rhythms could be explained by changing the circadian amplitude and increasing the drive to wake active neurons during wake. The close correspondence between the Phillips-Robinson model and the 2pm means that increasing the drive to wake promoting neurons is effectively the same as increasing the mean level of the upper threshold, see Supplementary Fig. S3.

### Polyphasic sleep in different species

Daan, Beersma & Borbély highlighted that the 2pm can explain many different sleep patterns [2], as also shown in Fig. 1 and Fig. 7. In particular they noted that when the thresholds are close together polyphasic sleep occurs. Similarly, Phillips, Chen and Robinson [40] argued that their mutual inhibition neuronal model could model observed patterns of sleep in different species. They showed that decreasing the homeostatic time constant for wake and sleep caused a shift from monophasic to polyphasic sleep and that increasing the mean drive to sleep active neurons increased the percentage of time spent asleep. In addition, Phillips et al. [41] argued that their mutual inhibition neuronal model could explain the fact that lesioning the SCN in squirrel monkeys removed the circadian rhythm of sleep and wake, as observed in [25].

The mathematical structure of the 2pm adds further insight to all three of these sets of observations. Reducing the gap between the thresholds reduces the natural period of the sleep-wake cycle *T*_nat_, see equation (1) and Fig. 6. When the natural period is short, periodic sleep-wake patterns which consist of multiple sleeps per day occur because of the underlying entrainment structure, as illustrated in Fig. 7.

The relationship between the neuronal model and the 2pm, means that the same underlying mathematical structure is at work to explain the observations in [40]. Reducing the homeostatic time constants reduces the natural period, resulting in polyphasic rhythms for small values of the time constants, see Fig. S3. Increasing the mean level of the drive to sleep active neurons is equivalent to lowering the mean level of the thresholds, which increases the percentage of the natural period in which time is spent asleep.

Similarly, the fact that setting the circadian amplitude to zero can result in a transition from monophasic to polyphasic sleep, as in [41], is also underpinned by the tongue structure. As can be seen in Fig. 7, at high circadian amplitudes, a wide band of natural sleep-wake periods are entrained to monophasic sleep. If the circadian amplitude is reduced to zero, then oscillations revert to the natural sleep-wake period. For the parameters chosen by Phillips et al, this resulted in a switch from monophasic sleep-wake patterns to patterns with a period of approximately 5h.

The mathematical structure in addition highlights that when the circadian amplitude is low, polyphasic sleep will be distributed evenly across the 24-hour day with little variance in duration of one sleep-wake cycle and the next. In contrast, when the circadian amplitude is high, sleep episodes will predominantly occur only at some circadian phases and the duration of sleep-wake cycles will vary with time-of-day, see [42] and also Supplementary Fig. S6.

An alternative hypothesis for the occurrence of polyphasic sleep in some species is that it is a consequence of noise resulting in random switching between wake and sleep. To understand the impact of noise in a two-process framework, it is perhaps useful to consider the potential formulation, see Supplementary Fig. S2. When sleep and wake both exist, switching between them depends on the amount of noise as compared with the height of the potential barrier between the two states. The height of the potential barrier depends on how close the state of the system is to the thresholds, so close to the upper threshold, sleep is the more likely state, and close to the lower threshold wake is the more likely state. Distance from the threshold depends on homeostatic sleep pressure and circadian rhythmicity. Noise alone, without sleep homeostasis or circadian rhythmicity would result in polyphasic sleep patterns which are randomly distributed across the day and night with wake and sleep bout duration normally distributed. Noise with circadian rhythmicity but without sleep homeostasis would result in polyphasic sleep patterns with sleep primarily occurring in one half of the circadian day. The predicted pattern with noise and both circadian rhythmicity and sleep homeostasis introduces further possibilities, as does the threshold shape.

The underlying mechanism for polyphasic sleep may be species specific. For example, in voles there is evidence for an additional ultradian rhythm generator [43]. In mice, there is evidence that the homeostatic time constants, as quantified by the rise and fall of SWA, are shorter than in humans. (Typical values for the homeostatic rise and fall time constants in humans have been reported as 18.2 h and 4.2 h respectively [2], whereas in mice the equivalent reported values are 8.3 h and 1.7 h [44]). An important point is that the two-process model provides a quantitative predictive framework to explore mechanistic questions.

### Transitions in early childhood

Developmental changes across early childhood result in changes from polyphasic sleep to monophasic sleep. The mathematical structure of the 2pm further suggests that this could be explained by physiological changes that lead to a gradual lengthening of the natural-sleep wake period rather than a change in the circadian process. In this progression, the theoretical structure predicts that periods of relatively regular sleep patterns followed by periods of transition, as the Arnold tongue structure is traversed [10]. For example in the shift from a pattern of three sleeps a day (a main nocturnal sleep, a morning nap, an afternoon nap) to two sleeps a day (a main nocturnal sleep and an afternoon nap) it is to be expected that there will be a progression when first there are only occasional days on which two sleeps occur through to a point where there are only occasional days in which three sleeps occur. The sequence follows a so-called Devil’s staircase, see Fig. 8. The particular example staircase shown in Fig. 8 was constructed by taking a slice across the tongue structure shown in Fig. 7 at the fixed value of the circadian amplitude *a* = 0.12. Evidence that the Devil’s staircase structure may help explain longitudinal data in infants has recently been provided by [45]. The authors considered data collected in four infants for between 179 and 546 days. They fitted a version of the Phillips-Robinson model [27] to investigate how time evolution of model parameters could best explain the data. Although there was considerable heterogeneity between the four infants, broad trends suggested that homeostatic parameters played a major role in the observed patterns of sleep over the first two years of human life. A second recent study, [46], fitted a (different) neuronal model to data describing the transition from napping to non-napping behaviour in children between the ages of two and five years. This second study also highlighted the fact that homeostatic parameters could describe the transition. In both cases, Arnold tongues and the associated Devil’s staircases provide the underpinning mathematical framework.

**Figure 8.**
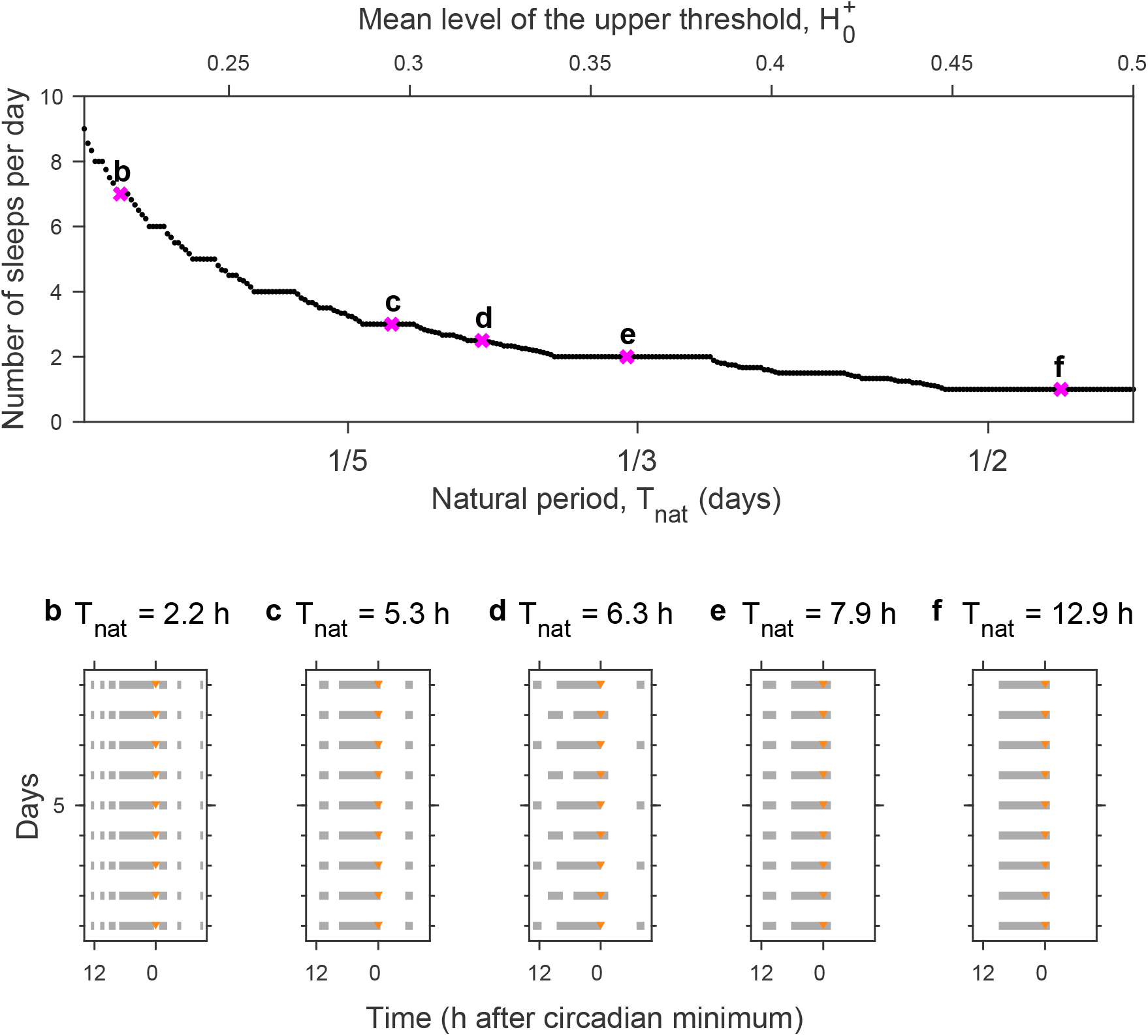
Devil’s staircase. As the natural period is increased for fixed circadian amplitude the number of sleeps per day follows a pattern known as a ‘Devil’s staircase’. Here this is illustrated for the fixed parameter values *χ*_*s*_ = 4.2 h, *χ*_*w*_ = 18.2 h, 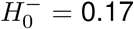, *µ* = 1 and *a* = 0.12 by varying the value of the upper threshold 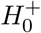 i.e. taking a horizontal line across Fig. 7a. Note the curvature of the tongues means that the plateau at, for example, *T*_nat_ = 1/3 corresponds to 2 sleeps a day. **b-f** Example raster plots for different values of 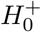.

### Timing of sleep relative to the 24 hour day: extending the two-process model to include the light-dark cycle

The 2pm models the internal phase relationship between sleep and circadian rhythmicity. Depending on the scenario being modelled, it is often implicitly assumed that the endogenous circadian period is entrained to 24-h. Timing relative to the 24-h day has been modelled with additional assumptions on the clock time of the circadian minimum, for example assuming that it occurs at 04:00, as in Fig. 1a. However, the time of the circadian minimum varies both within and between people. To fully model timing relative to the 24-h day requires the inclusion of the synchronisers (‘zeitgebers’ i.e. ‘time giver’) of the circadian pacemaker.

Extensive laboratory experiments have shown that light is the primary zeitgeber of the human circadian clock. Many experiments have quantified the effect of light, and mathematical equations that quantitatively describe the phase and dose response of the human circadian pacemaker to light have been derived, originally by Kronauer [47, 28] and subsequently modified by others, for example [48, 49]. The conceptual and mathematical understanding is that the light-circadian system is a forced oscillator system in which the intrinsic circadian oscillation is entrained by the light-dark cycle. The circadian-light forced oscillator system, in common with other nonlinear forced oscillator systems, including the sleep-circadian forced oscillator system, has an underlying mathematical structure in which there are entrainment regions that form tongue-like regions in the (natural period)–(forcing strength) parameter space [28, 50, 27]. If the light-dark cycle is only a weak zeitgeber i.e. there is insufficient contrast between light that is available during the day and light that is available during the night, then only a small range of intrinsic circadian periods close to 24 hours can be entrained. The stronger the forcing, the larger the range of circadian periods which can be entrained. Consequently, the entrainment regions form tongue-like structures. Typically of human relevance, the focus is usually on the (1,1) tongue in which one 24 hour light-dark cycle results in one 24 hour entrained circadian oscillation, but in principle other tongues exist. Analogous to internal desynchrony for the sleep-circadian oscillator system, desynchrony with respect the 24 hour day occurs when the light-dark cycle is too weak or the intrinsic period is too far from 24 hours. Light-circadian models have been suggested as a method for evaluating circadian phase in humans [51, 52, 53].

To capture sleep and circadian phenotypes requires combining both light-circadian and circadian-sleep models. In this hierarchical entrainment framework, light entrains the circadian rhythm, and the circadian rhythm entrains the sleep-wake cycle. However, crucially, there is feedback between sleep-wake patterns and the timing of light (or other zeitgebers). Unique to humans, access to electric light enables us to turn light on after dusk. The timing of the circadian entraining signal is then dictated by the time at which we turn the lights off and the time at which we wake up in the morning. The consequent feedback between sleep behaviour and the timing of the entraining signal has a ‘destablizing’ effect which is critical for understanding human chronotypes [27].

Early models that modelled the light-circadian-sleep homeostasis system include work by Phillips, Chen & Robinson [54], who extended the neuronal Phillips-Robinson model Using the neuronal framework, plausible physiological reasons to explain changes in sleep duration and timing across the lifespan have been proposed [55]. The light-circadian-sleep neuronal framework has been further adapted and used to model the effects of self-selection of light on sleep timing and the effects of social constraints which restrict the opportunities to sleep [27]. The version of the neuronal light-circadian-sleep model in [27] has been shown to capture individual sleep phenotypes [56] and used for scenario testing of, for example, the effect on social jetlag of moving school start times versus changing the light environment. Studying sleep in those living with schizophrenia, it has been suggested that a plausible reason for long sleep duration is a reduced wake propensity (reduced level of the mean thresholds in a two-process representation) [56]. Furthermore, fitting to individuals, highlighted that observed seasonal desynchrony could be plausibly explained by reduced light exposure in the winter months [56].

Inspired by the neuronal sleep-circadian-light neuronal framework and the close link between neuronal models and the 2pm, a homeostatic-circadian-light (HCL) model has been proposed, as represented schematically in Fig. 9. The HCL model combines the graphical simplicity of the original 2pm with a light-circadian entrainment process and social constraints [21]. At the same time, the shape of the thresholds was fitted to data for the circadian drive for wakefulness, as reported in [4]. The updated shape of the upper threshold better captures well-established features of sleep-wake regulation such as the wake maintenance zone [57, 58]. It has been shown that combining the HCL model with data, personalised models which capture individual average sleep duration and timing can be constructed. In [21], the model was used to develop a method to separate out the relative contributions of physiology and environmental light in determining sleep timing. These personalised models could be used to design personalised light interventions to adjust sleep timing [56, 21].

**Figure 9.**
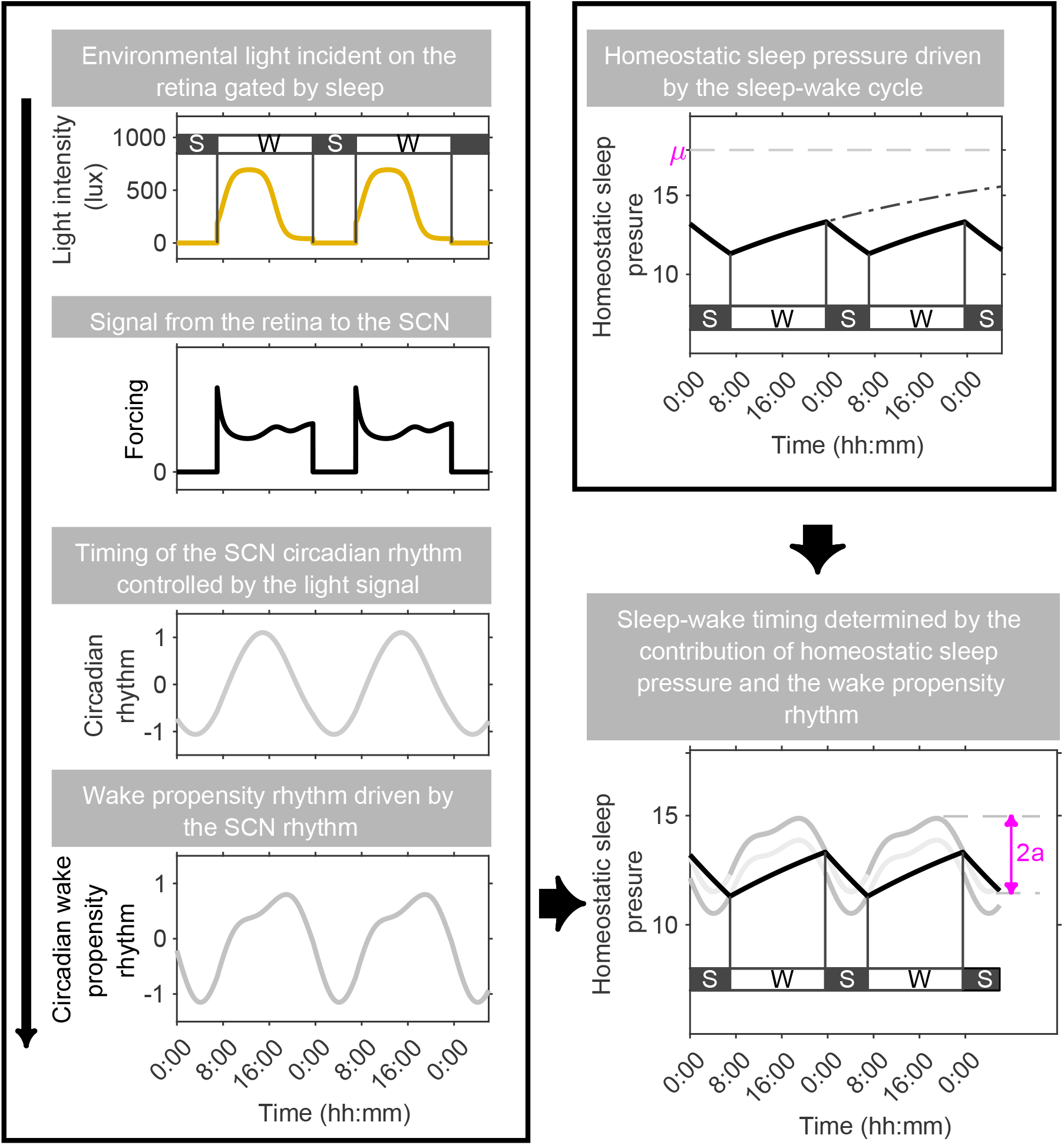
The homeostatic-circadian-light (HCL) model. Adapted from [21]. In the HCL model, light, gated by sleep, is fed into a model which captures the phase and dose response of light on the circadian pacemaker [48]. The circadian pacemaker then results in a wake propensity rhythm which, when combined with sleep homeostasis, results in patterns of sleep and wake.

## Discussion

One enduring appeal of the 2pm is its graphical simplicity which gives an intuitive understanding of how sleep duration depends on time-of-circadian-day. Aside from its use to frame experimental results, all current models of fatigue risk management are to a greater or lesser extent based on the principles of sleep homeostasis and circadian rhythmicity [59].

The 2pm is often represented as a dichotomy of two separate processes, sleep homeostasis and circadian rhythmicity, which has prompted the ‘separation’ of the processes in forced desynchrony protocols [60, 61]. However, interestingly, the 2pm is all about interactions. It is the interaction of the two processes that explains, for example, the duration of recovery sleep following sleep deprivation. It is the interaction that means that polyphasic sleep occurs at consolidated circadian phases when the circadian amplitude is high [42].

Indeed, it is also not so easy to separate out what are ‘homeostatic’ and ‘circadian’ parameters [62]. Specifically, of the eight parameters in the 2pm, the rate constants *χ*_*w*_ and *χ*_*s*_ which represent the dissipation rate of homeostatic sleep pressure during wake and sleep respectively may be described as homeostatic parameters. Similarly, one may argue that *µ*_*w*_ and *µ*_*s*_ which represent the rate of increase of homeostatic sleep pressure during wake and sleep respectively are homeostatic parameters. The circadian amplitude *a* is evidently a ‘circadian’ parameter. However, this raises a question about the two threshold parameters, the mean levels of the upper and lower thresholds, 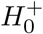 and 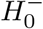 respectively, which have often been considered circadian parameters [2]. Relating the 2pm to neuronal mutual inhibition models suggests that these thresholds may relate to properties of the neuronal mutual inhibition structure of sleep-wake regulation and inputs to wake promoting and sleep promoting regions — and may therefore be neither circadian nor homeostatic parameters.

Linking the 2pm to neuronal mutual inhibition model also raises the question as to what is ‘the two-process model’, is it a specific set of parameters? Conceptualising the 2pm as instead a hysteretic loop suggests that there is no physiological reason for homeostatic sleep pressure to remain ‘between’ the thresholds. It also highlights the fact that the 2pm may alternatively be posed as a double-well potential problem in which the time dependent drives change the probability of states of wake and sleep occurring.

The 2pm has a deep mathematical structure that is inherent to nonlinear forced oscillator systems, as highlighted in Fig. 7 [19]. Understanding the mathematical entrainment structure establishes what determines different sleep-wake patterns in the 2pm. For sufficiently small circadian parameters (where ‘sufficiently small’ has a precise definition in terms of the parameters [19]) stable solutions are unique, so the only way that different patterns can be observed is by changing parameters. For high circadian amplitudes, multiple solutions may be possible signifying that different patterns could occur as a change of state rather than a change of parameters. It is also likely that there are regions of parameter space in which monophasic sleep-wake patterns occur but with alternating long and short sleep duration.

A further open question is the optimal parameters to describe sleep-wake regulation in humans, cf, Fig. 1b and Fig. 3g. Slow wave activity (SWA) is viewed as a marker of sleep homeostasis [1, 63, 7], but it may or may not be that the time constants found by fitting to the timecourse of SWA are the same as the time constants of a homeostatic sleep process which determines sleep timing. The time constants used in the Phillips-Robinson model are typically much longer (∼45 hours) yet can still replicate similar patterns of sleep. The use of longer time constants may be appropriate when, for example, simulating the effects of acute and partial sleep deprivation on performance and sleepiness [64, 65].

Understanding the mathematical structure can also challenge our pre-conceptions on what causes phenotypes or phenotypic change in humans. For example, changes from polyphasic to monophasic sleep may not primarily be a consequences of maturation of the circadian system [10, 45]. Changes in sleep timing as we age may be a consequence of changes in circadian amplitude and / or sleep homeostasis and not changes in circadian period [55]. Differences in sleep duration may be a consequence of differences in the drive to wake or the drive to sleep [56], and not because of changes in homeostatic time constants. Desynchrony with the 24-hour day may not be due to clock-malfunction but due to insufficient zeitgeber strength [56].

Finally in Fig. 1 of their paper, Daan, Beersma & Borbély set out a scheme for their model [2]. In this scheme, the model included the effect of the light-dark cycle to synchronise the circadian oscillator and conscious decisions affecting the thresholds as well as the interactions of sleep homeostasis and circadian rhythmicity. With extensions of the 2pm to include input from the light-dark cycle we are perhaps now somewhat closer to this vision [21]. Even though the 2pm does not capture basic features of sleep-wake regulation such as the REM—non-REM cycle, there remains much promise with a ‘four-process’ framework which includes sleep homeostasis, circadian rhythmicity, light and dynamic thresholds which encompass acute and chronic effects of stimulants [66, 67].

Most of all, it is of value to recognise that the 2pm and its extensions are not only conceptual frameworks, but quantitative mathematical models which can be fitted and challenged by data. Testing models in this quantitative way brings the opportunity to evaluate whether or not they can really explain the available data and what is missing. This ultimately may inform understanding of the wide variety of sleep phenotypes, sleep-wake disorders and the development of countermeasures for situations in which the sleep-wake and circadian system is challenged by social constraints.

## Methods

The two-process model (2pm) of sleep regulation posits that there is a homeostatic sleep pressure *H*(*t*) which increases during wake and dissipates during sleep. During wake, it is typically modelled by

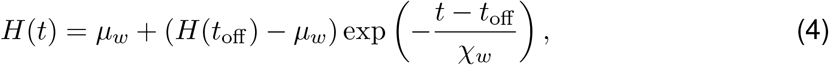

where *H*(*t*_off_) *< µ*_*w*_. While during sleep, homeostatic sleep pressure is typically modelled by

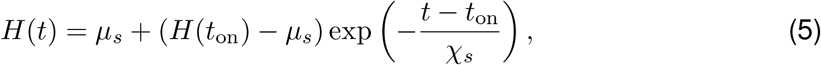

where *H*(*t*_on_) *> µ*_*s*_, *t* is time, *t*_on_ and *t*_off_ are the times of sleep onset and offset respectively. The parameters *µ*_*w*_ and *µ*_*s*_ are the upper and lower asymptotes respectively. The parameters *χ*_*w*_ and *χ*_*s*_ are time constants for the homeostatic sleep pressure during wake and sleep respectively. The condition *H*(*t*_off_) *< µ*_*w*_ for equation (4) means that homeostatic sleep pressure increases during wake, approaching the upper asymptote value *µ*_*w*_ as *t* → ∞. During wake, the rise of homeostatic sleep pressure is governed by both *µ*_*w*_ and *χ*_*w*_. The condition *H*(*t*_on_) *> µ*_*s*_ for equation (5) means that homeostatic sleep pressure decreases during sleep, approaching the lower asymptote value *µ*_*s*_ as *t* → ∞. During sleep, the decay rate of homeostatic sleep pressure is governed by both *µ*_*s*_ and *χ*_*s*_.

When equations (4) and (5) are used to estimate the rate parameters *χ*_*s*_ and *χ*_*w*_ from data, an essential step is an estimate of *µ*_*w*_ and *µ*_*s*_.

In differential equation form the 2pm is

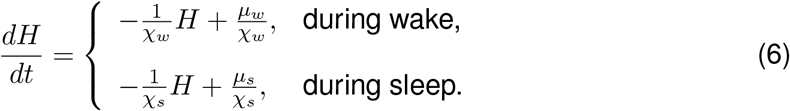

These equations mean that the net rate of change of homeostatic sleep pressure is a balance between a loss term and a gain term. Loss, as represented by the first term on the righthand sides, is proportional to the instantaneous value of homeostatic sleep pressure and occurs at a rate of 1*/χ*_*w*_ during wake and 1*/χ*_*s*_ during sleep. Gain is assumed to occur at a fixed rate of *µ*_*w*_*/χ*_*w*_ during wake and *µ*_*s*_*/χ*_*s*_ during sleep.

Now the 2pm, as framed by Daan, Beersma & Borbély [2] considers that sleep-wake cycles are a result of switching from wake to sleep and from sleep to wake which occurs at thresholds which are periodically modulated to mimic the effect of circadian rhythmicity. Hence, switching from wake to sleep occurs at *H*(*t*) = *H*^+^(*t*) where

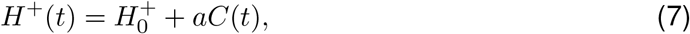

and switching from sleep to wake occurs at *H*(*t*) = *H*^*−*^(*t*) where

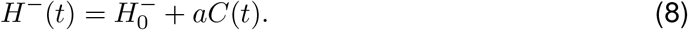

The constants 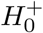 and 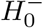 are the mean levels of upper and lower thresholds and the wave form of the time dependent circadian modulation *C*(*t*) represents the circadian rhythm of wake propensity. In its simplest form *C*(*t*) is often taken to be sinusoidal, *C*(*t*) = cos *ωt*, where *ω* is the angular frequency of the circadian oscillator i.e. the circadian period is 2*π/ω*.

The full model thus has eight parameters (*χ*_*w*_, *χ*_*s*_, *µ*_*w*_, *µ*_*s*_, 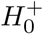, 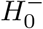, *a* and *ω*).

Daan, Beersma & Borbély et al [2] realised that, from the point of view of understanding the behaviour of this model shifting and rescaling the equations could effectively remove two of these eight parameters. Specifically, by shifting 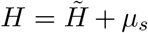, equations (6) become

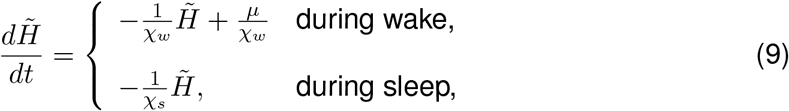

where *µ* = *µ*_*w*_ − *µ*_*s*_. Then during wake,

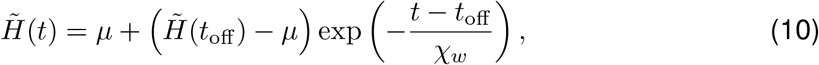

and during sleep

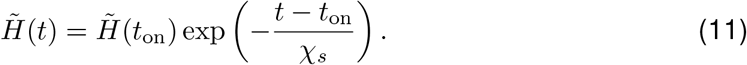

By shifting and scaling i.e. setting, 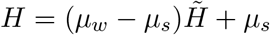, equations (6) become

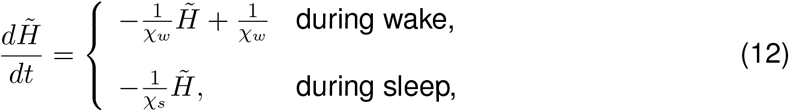

Then during wake,

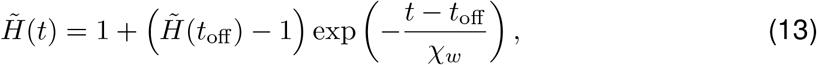

and during sleep

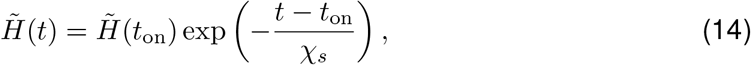

where now switching occurs at thresholds

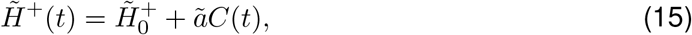

and switching from sleep to wake occurs at 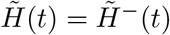 where

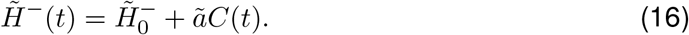

Where 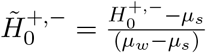 and and 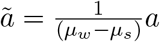.

In the absence of circadian rhythmicity the duration of a sleep episode is calculated as follows. Suppose sleep starts at *t* = *t*_on_ which occurs when homeostatic sleep pressure is at the upper threshold, i.e. 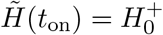. Sleep ends at *t* = *t*_off_ when homeostatic sleep pressure reaches the lower threshold, i.e. 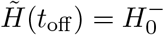 Then substituting into equation (11) which describes the time evolution of homeostatic sleep pressure during sleep gives,

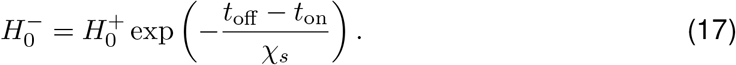

Now, the duration of sleep *T*_sleep_ is by definition *t*_off_ − *t*_on_, so

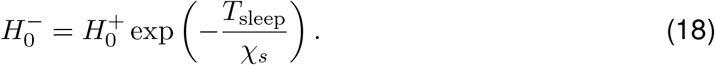

Re-arranging

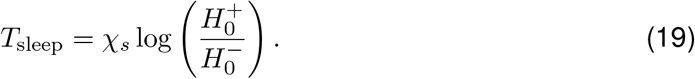

Similarly, the duration of a wake episode can be calculated as follows. Suppose wake starts starts at *t* = *T*_off_ which occurs when homeostatic sleep pressure is at the lower threshold, i.e. 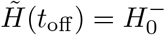. Wake ends at *t* = *t*_on_ when homeostatic sleep pressure reaches the upper threshold, i.e. 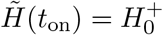. Then substituting into equation (10) which describes the time evolution of homeostatic sleep pressure during wake gives,

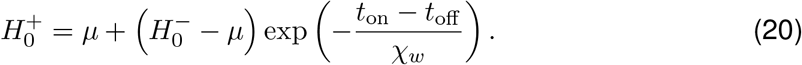

Now, the duration of wake T_wake_ is by definition *t*_on_ -*t*_off_, so

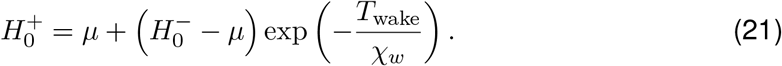

On re-arranging, this leads to

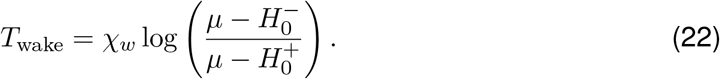

Simulations of the 2pm were carried out in MATLAB [68].

## Data availability

Data reported in this article will be made available on github on acceptance of this article.

## Code availability

Code to simulate the 2pm will be made available on github on acceptance of this article.

## Acknowledgments

The authors are supported by the UK Dementia Research Institute [award number UKDRI-7206] through UK DRI Ltd, principally funded by the UK Medical Research Council, and additional funding partner Alzheimer’s Society and also by the National Institute for Health Research (NIHR) Oxford Health Biomedical Research Centre (BRC), (NIHR203316). The views expressed are those of the authors and not necessarily those of the NHS, the NIHR or the Department of Health. This paper is based on a long-standing collaboration and many discussions between the authors, and contributions from others including Andrew Phillips and Gianne Derks. We thank Stanisław Biber for feedback on the paper. This paper is in honour of and to celebrate the 51st anniversary of the Daan, Beersma and Borbély 1984 paper.

## Author contributions

ACS produced the first draft of the manuscript. DJD and ACS reviewed and edited the manuscript.

