## Supplementary Figures for "The complexity and commonness of the two-process model of sleep regulation from a mathematical perspective"

### Supplementary Material

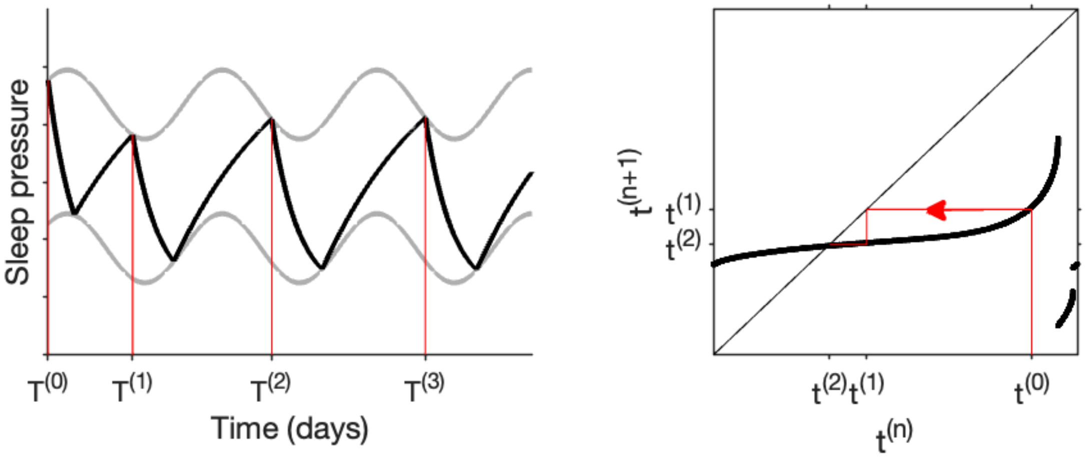


Figure S1: **Circle map description of the two-process model.** Simulating the two-process model generates a sequence of sleep onset times, $T^{(0)}, T^{(1)}, T^{(2)} \ldots$, as shown in the lefthand panel. In the example shown, $T^{(0)} = 0.875, T^{(1)} = 1.421, T^{(2)} = 2.319 \ldots$, where time is in days and is measured from the circadian maximum. In other words, the two-process model (2pm) is a rule that takes a sleep onset time $T^{(n)}$ and predicts the next sleep onset time $T^{(n+1)}$. However, time is 'circular', as reflected by the fact that the thresholds are periodic. So, considering time-of-day by taking modulo 1, the 2pm generates a sequence of time-of-day values $t^{(0)}, t^{(1)}, t^{(2)} \ldots$, where in the example shown, $t^{(0)} = 0.875, t^{(1)} = 0.421, t^{(2)} = 0.319 \ldots$. The dynamics of the two-process model can then be understood by considering all possible values of $t^{(n)} \in [0, 1)$ and evaluating the resulting $t^{(n+1)}$. The result is the circle map shown in the lefthand panel. Superimposed on the circle map is the sequence of times generated by the starting condition $t^{(0)} = 0.875$ in a process known as 'cobwebbing'. In this example, it can be seen that all solutions converge to a stable periodic solution with sleep onset at $t \approx 0.311$. Circadian audiences may be familiar with 'phase transition curves', which are also examples of circle maps.

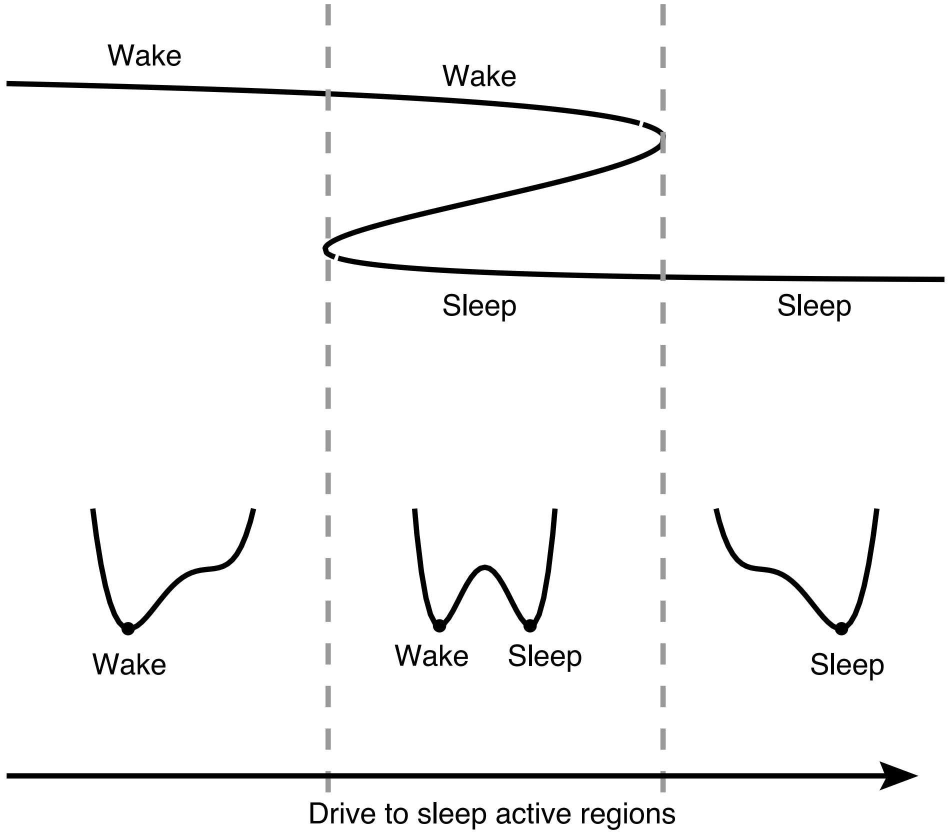


Figure S2: **Potential formulation.** The Phillips-Robinson model and, by extension, the two-process model may alternatively be viewed as a system in which there are two states, sleep and wake. As sleep homeostasis and circadian rhythmicity change, the relative depth of the potential wells for each state changes, changing the likelihood that one or other state exists.

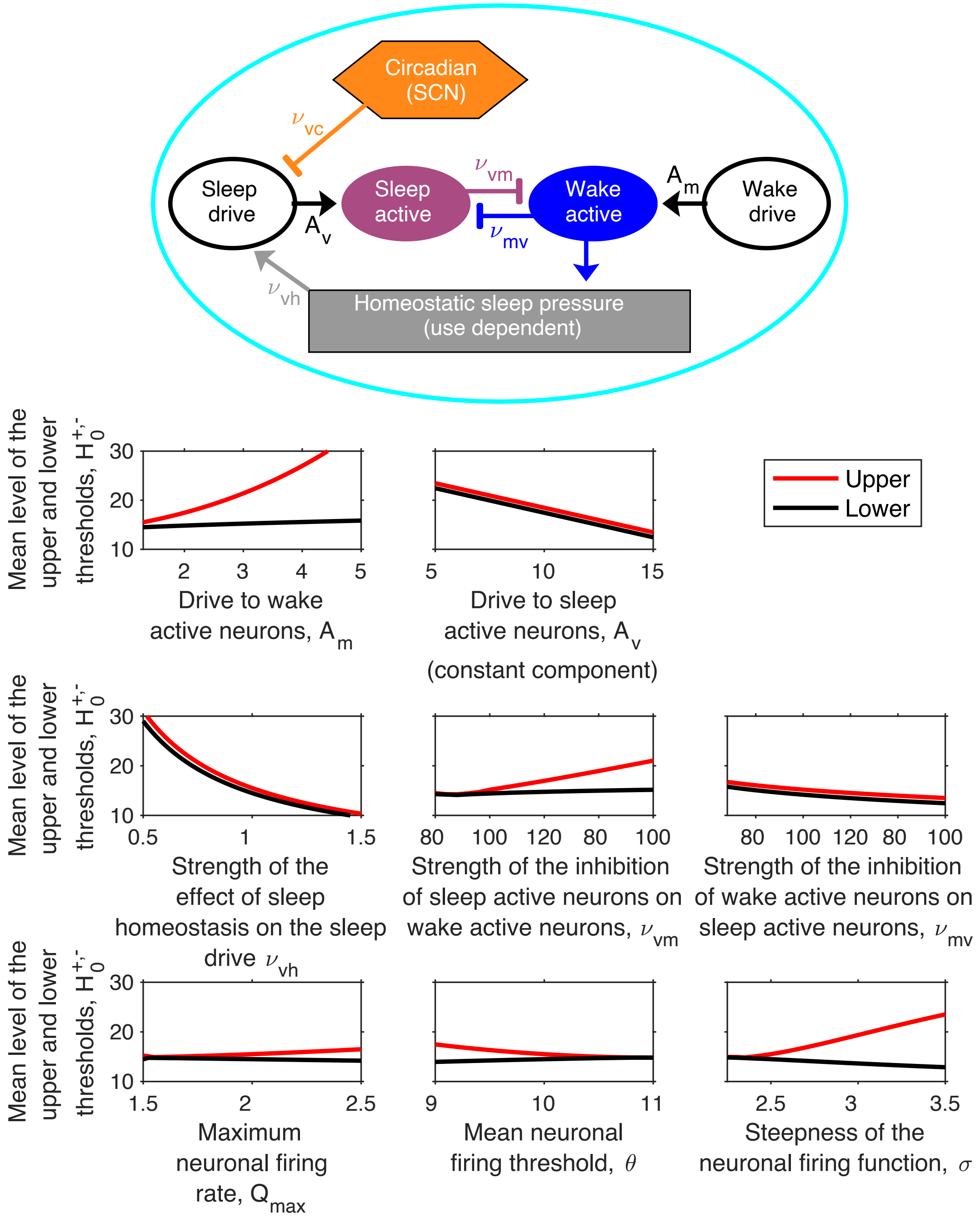


Figure S3: **Relation of the thresholds to neuronal parameters.** The mean level of the upper and lower thresholds in the two-process model for different values of the parameters in the Phillips-Robinson neuronal mutual inhibition model. Values are calculated from [10]. We also note that, as framed in the neuronal models, the mean level of the thresholds is dependent on the circadian amplitude. This is essentially a modelling choice and relates to whether, physiologically, increasing circadian amplitude results in a larger amplitude but no change in the mean value or, alternatively, increasing circadian amplitude results in a larger amplitude with no change to the minimum value. To help visualise where the parameters $A_m, A_v, \nu_{vh}, \nu_{vm}$ and $\nu_{mv}$ appear in the Phillips-Robinson model they are shown on the relevant arrows in the network diagram in the top panel. The parameters $Q_{\max}, \theta$ and $\sigma$ relate to the neuronal firing function and therefore contribute to the inhibition of wake promoting neurons by sleep promoting neurons, the inhibition of sleep promoting neurons and the impact of the firing of wake promoting neurons on homeostatic sleep pressure.

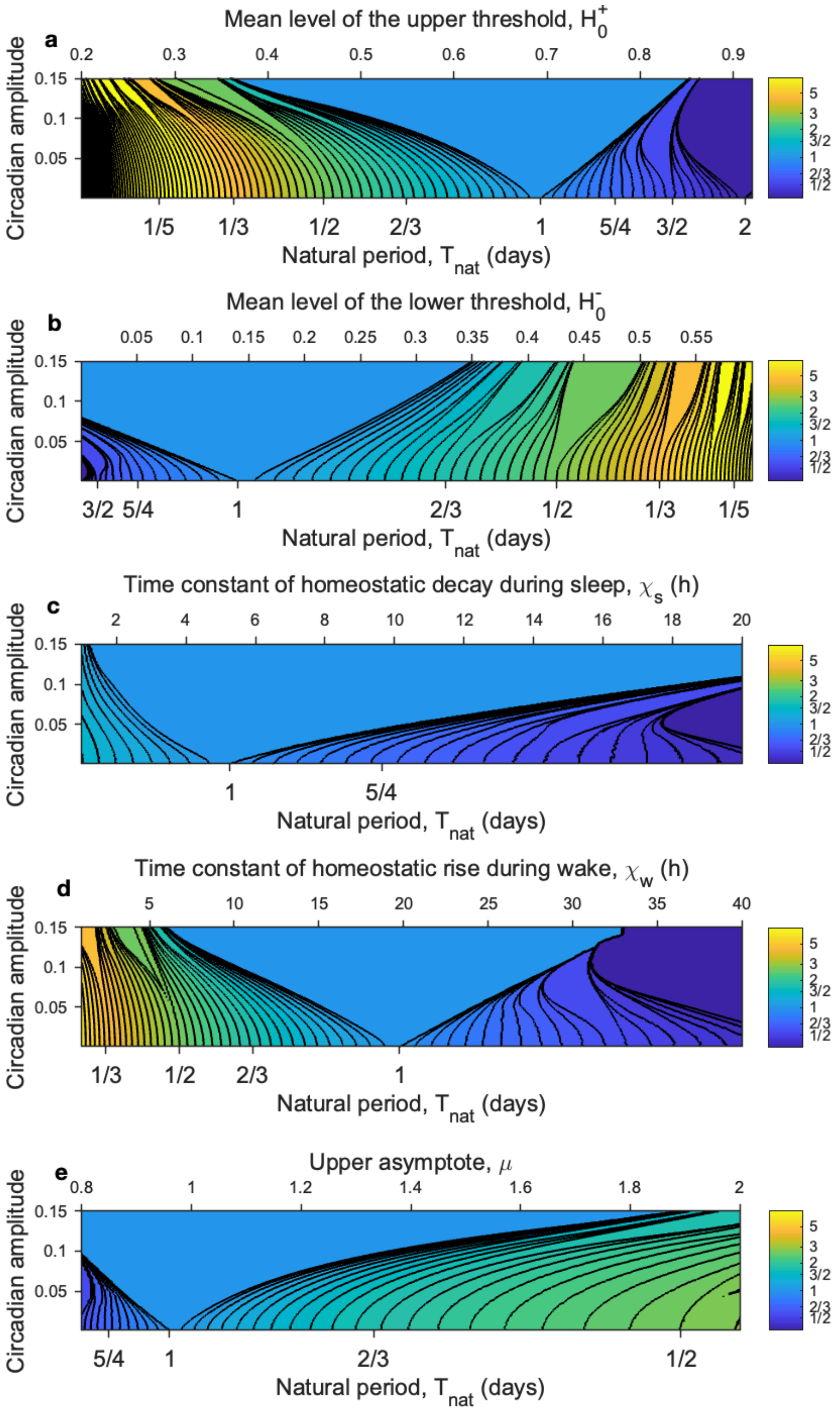


Figure S4: **Entrainment of the sleep-wake oscillator by the circadian oscillator: Arnold tongues.** Each panel shows the result of simulating the two-process model for approximately 100,000 different parameter values. In each case the number of sleeps per day as a function of the circadian amplitude and one of the other parameters of the model is shown. The default parameters are: $\chi_s = 4.2$ h, $\chi_w = 18.2$ h, $H_0^- = 0.17$, $H_0^+ = 0.67$, $\mu = 1$. Regions are shaded according to the number of sleeps per day. The black lines are contour lines. Both the shading and the contours are on a logarithmic scale (base 2). A similar tongue structure is shown in each case, but because of the relationship between the natural period and the parameters, each plot is squashed or stretched in different ways. Depending on the range of parameters used, panels cover a different range in natural periods $T_{\mathrm{nat}}$.

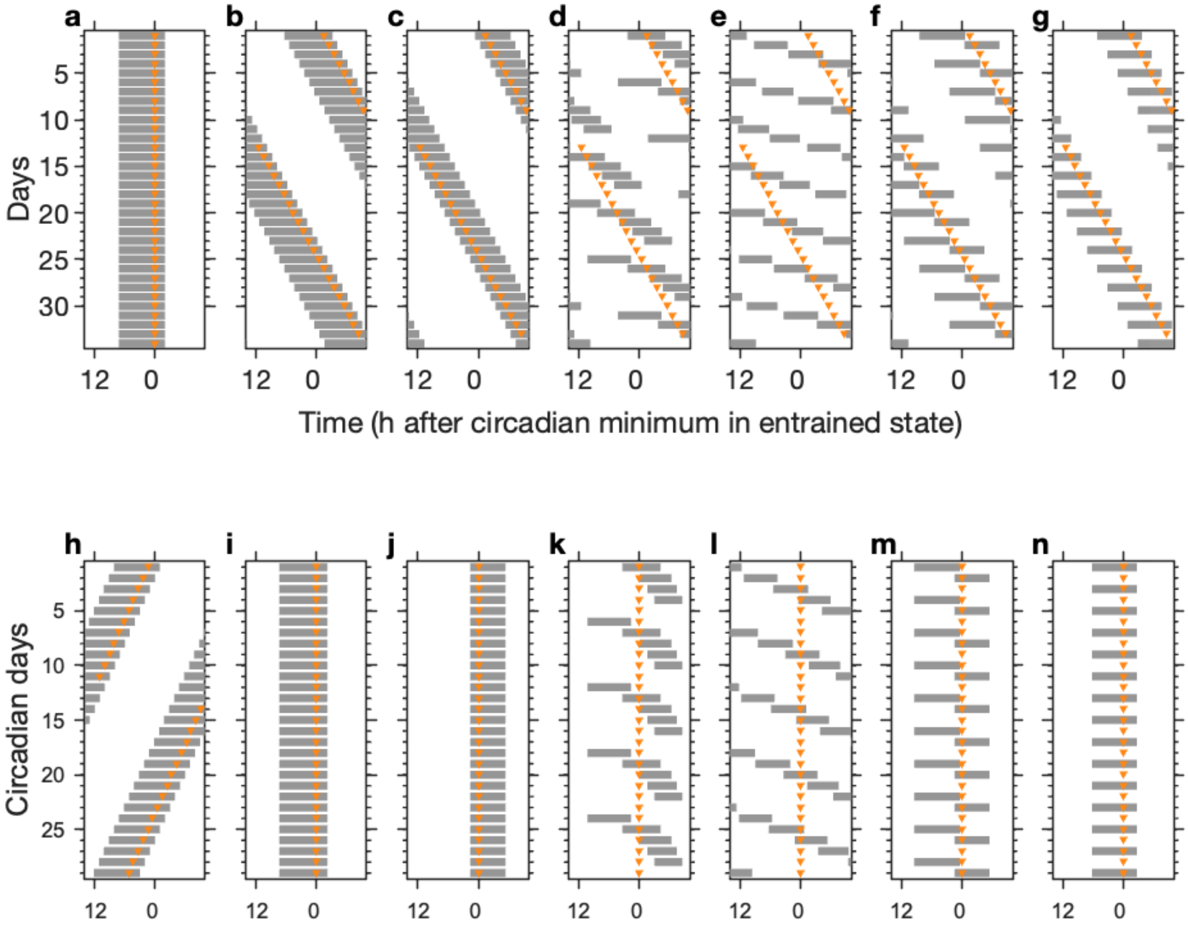


Figure S5: **Simulating patterns of sleep and circadian rhythmicity in time isolation experiments.** Horizontal grey bars show the timing of sleep on successive 24 hour days. The orange triangles mark the circadian minimum. For all panels, $\chi_s = 4.2$ h, $\chi_w = 18.2$ h, $H_0^- = 0.17$, $\mu = 1$. **a** Monophasic sleep entrained to 24 h. $T_f = 24$ h, $H_0^+ = 0.55$, $a = 0.12$. **b** Monophasic sleep entrained to 25 h. $T_f = 25$ h, $H_0^+ = 0.55$, $a = 0.12$. **c** Monophasic sleep entrained to 25 h, but with the circadian minimum at the beginning of the night. $T_f = 25$ h, $H_0^+ = 0.78$, $a = 0.12$. **d** Internal desynchrony with five sleeps every six circadian days. $T_f = 25$ h, $H_0^+ = 0.7925$, $a = 0.07$. **e** Internal desynchrony with sleep-wake periods evenly distributed across the circadian day. $T_f = 25$ h, $H_0^+ = 0.78$, $a = 0.02$. **f** Three sleeps every two circadian days. $T_f = 25$ h, $H_0^+ = 0.82$, $a = 0.07$. **g** Bicircadian rhythms with one sleep every two circadian days. $T_f = 25$ h, $H_0^+ = 0.90$, $a = 0.07$. bf h - n show the same data as **a - g** but wrapped at 25 h rather than 24 h.

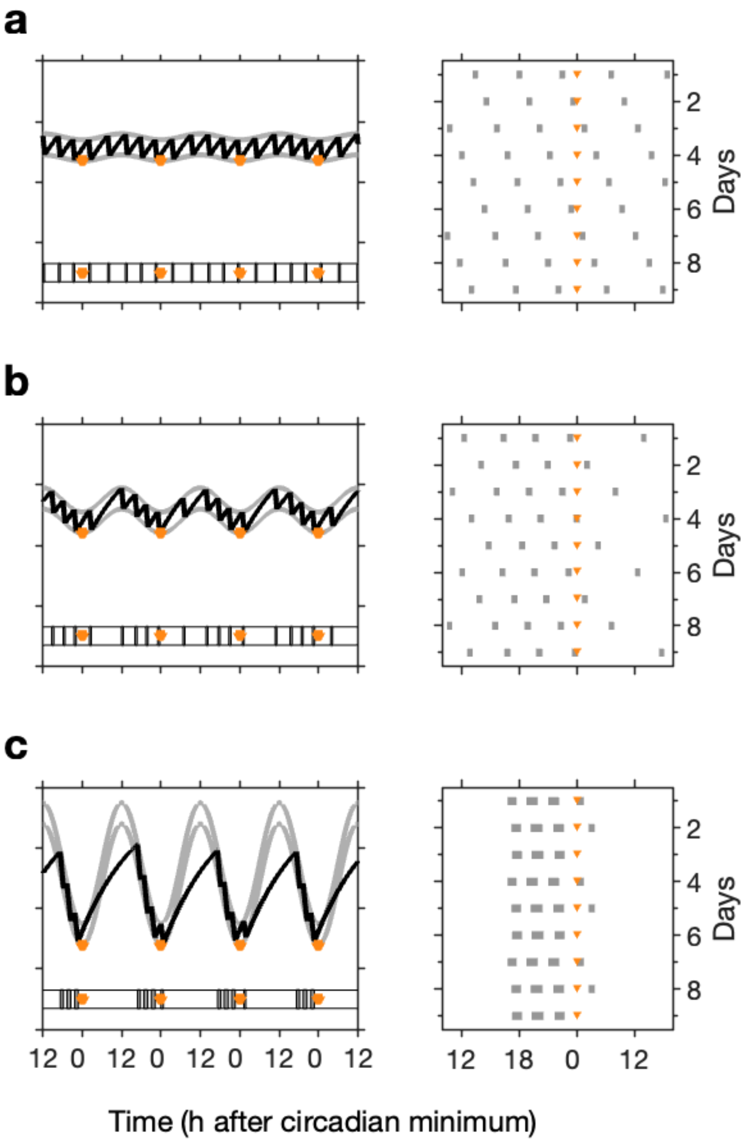


Figure S6: **Distribution of polyphasic sleep patterns across the 24 hour day depends on circadian amplitude**. Simulations of the two-process model for $\chi_s = 6$ h, $\chi_w = 18$ h, $H_0^+ = 0.75$, $H_0^- = 0.68$, $\mu = 1$ and for varying circadian amplitude $a$. For low circadian amplitudes, sleep is evenly distributed across the day. Increasing circadian amplitude has the effect of consolidating sleep so that it primarily occurs at some circadian phases. **a** $a$ =0.01. **b** $a$ =0.04. **c** $a = 0.2$. In each panel, to the left is shown the two-process model simulation with the horizontal bars showing the timing of sleep. To the right is shown a raster plot showing the distribution of sleep over 10 successive days. The orange triangles mark the positions of the circadian minima.
